# Mechanistic insight into the regulation of virulence factor type 1 pili in pathogenic *E. coli* by rhomboid protease GlpG

**DOI:** 10.1101/2025.09.09.675128

**Authors:** Jimmy Lu, Elena Arutyunova, Bridgette Hartley, Jonas Wong, Hyunjae Chung, Lanny Buchan, Heather Armstrong, Sadhna Phanse, Howard S. Young, Eytan Wine, Mohan Babu, Justin Chun, Wael Elhenawy, Olivier Julien, M. Joanne Lemieux

## Abstract

With the rise in antimicrobial resistance, understanding the virulence factors utilized by pathogenic *E. coli* is essential for the development of alternative therapeutics. While previous work has shown that disruption of the *E. coli* rhomboid protease gene *glpG* leads to defects in bacterial colonization, here we provide mechanistic insight into the loss of fitness. We show GlpG is essential for the assembly of type 1 pili, a virulence factor required for the colonization of eukaryotic cells. Since pili are critical for biofilm formation and bacterial persistence, the absence of GlpG proteolytic activity reduces the production of biofilm. Working towards new potential antimicrobial targets for treating infections, we show that biofilm formation is hampered by GlpG inhibition. Our data demonstrates that GlpG plays a key role in protein quality control of type 1 pili and alters the paradigm for GlpG proteolysis, previously implicated in the cleavage of only membrane embedded substrates.

## Introduction

*Escherichia coli* is a diverse bacterial species with the ability to colonize and persist in various niches within animal hosts, often forming a symbiotic relationship by providing nutrients, key signals for homeostatic regulation, and protection against foreign pathogens (Nataro James and Kaper James, 1998; Tenaillon et al., 2010; van Elsas et al., 2010). Within the gastrointestinal tract, resident *E. coli* are typically commensal, however these strains can take on a pathogenic course, causing disease outside the gut through combinations of acquired virulence factors such as adhesins, secreted toxins or lipopolysaccharides (Croxen et al., 2013; Kaper et al., 2004; Vila et al., 2016). Pathogenic strains are broadly categorized as either intestinal pathogenic *E. coli*, causing infection within the gastrointestinal tract, or extraintestinal pathogenic *E. coli* (ExPEC) as they have retained the ability to survive in the gut, but also wield virulence factors with the capacity to disseminate and colonize other sites including the blood, brain, and urinary tract (Croxen and Finlay, 2010; Pakbin et al., 2021; Sarowska et al., 2019). In addition, there are three unclassified pathogenic groups: adherent-invasive *E. coli* (AIEC) involved in Crohn’s disease, necrotoxic *E. coli* (NTEC) which is associated with disease in humans and other mammals, and cell-detaching *E. coli* (CDEC) that involved in hemolysin production. Among ExPEC, the uropathogenic *E. coli* (UPEC) pathotype accounts for 80-90% of all urinary tract infections (UTI), one of the most common bacterial infections globally (Marrs et al., 2005; Subashchandrabose and Mobley, 2015; Terlizzi et al., 2017). About 50% of women in the United States are expected to have at least one UTI in their lifetime with half experiencing a recurrent infection within 12 months of the initial episode (Rozwadowski and Gawel, 2022; Salvatore et al., 2011). The global death rate due to UTIs has increased by 140% from 1990 to 2019, largely attributed to the rise of antimicrobial resistance to classes of commonly prescribed antibiotics such as β-lactams, cephalosporins and fluoroquinolones, ultimately making UPEC infections more challenging to treat (Darby et al., 2023; Flores-Mireles et al., 2015; Gupta et al., 2011; Mazzariol et al., 2017; Murray et al., 2022; Yang and Larson, 1998). Consequently, this places an urgent need for the development of alternate treatment strategies to combat these infections. However, in order to explore alternative therapeutics, a better understanding of bacterial virulence factors regulation is required.

Virulence factors play a crucial role in bacterial survival and persistence, largely through biofilm initiation and formation. Biofilms are surface-attached communities of bacteria embedded in a matrix of biopolymers such as proteins and polysaccharides, and are one of the critical properties responsible for most chronic infections by UPEC (Costerton et al., 1999). Pili and flagella, along with other adhesins and extracellular factors such as toxins, siderophores, enzymes, and polysaccharide coatings, are among the main virulence factors necessary for initial adhesion and colonization of host mucosal surfaces (Belas, 2014; Pratt and Kolter, 1998; Tabibian et al., 2008). Once formed, biofilms are remarkably difficult to eradicate as these communities are more resistant to antimicrobial agents, surviving transient antibiotic treatment and regrowing when antibiotic is withdrawn, leading to infection persistence and recurrence (Ito et al., 2009; Mah, 2012; Sharma et al., 2016).

The expression, assembly and turnover of virulence factors reflect a prokaryote’s ability to disseminate and infect a host, but how the proteins involved regulate these processes is not completely understood. The rhomboid protease superfamily are intramembrane proteases that are ubiquitously expressed across all kingdoms of life, with the eukaryotic members implicated in various functions such as mitophagy, growth factor signaling, amyloid precursor protein processing and more (Düsterhöft et al., 2017; Freeman, 2008; Lysyk et al., 2020; Paschkowsky et al., 2016; Wunderle et al., 2016). Meanwhile, the roles of prokaryotic rhomboid proteases have been linked to functions such as quorum sensing with AarA in *Providencia stuartii*, biofilm formation with Rho1/2 in *Haloferax volcanii*, ion homeostasis with YqgP in *Bacillus subtilis* and protein quality control with GlpG in *Shigella sonnei* (Began et al., 2020; Costa et al., 2025; Liu et al., 2020; Stevenson et al., 2007). GlpG is a conserved and widely represented member of the rhomboid protease superfamily located in the plasma membrane of all known strains of *E. coli* as well as other prokaryotes (Koonin et al., 2003). Moreover, it has been shown that the *glpG* gene is essential for ExPEC persistence in the murine intestine and by knocking-out *glpG*, bacterial colonization is significantly reduced, however by an unknown mechanism (Russell et al., 2017). Additionally, while GlpG in *S. sonnei* targets and cleaves orphan proteins to prevent aggregation in the cell, a potential association to bacterial pathogenicity remains to be shown (Liu et al., 2020).

Here, we provide a mechanistic insight into the link between bacterial colonization and proteostasis in pathogenic *E. coli*. Using proteomics, we sought to reveal physiological binding partners of GlpG and identified type 1 pili subunit, FimA, as well as periplasmic serine endoproteases DegQ and DegP (Kolmar et al., 1996). We demonstrate that GlpG proteolytic activity is essential for type 1 pili formation in pathogenic pathotype UPEC, affecting the ability of *E. coli* to colonize and infect host cells. As adherence is essential for the initiation of biofilm formation, we show that the absence of GlpG catalysis, as well as GlpG inhibition, significantly reduces biofilm production. Our data demonstrates that GlpG regulates FimA protein quality control and turnover by cleaving FimA aggregates during type 1 pili formation. Moreover, we demonstrate that GlpG associates and works in concert with periplasmic housekeeping proteases DegQ and DegP. We reveal that these proteases are also involved in cleaving aggregated or misfolded FimA subunits, assisting in the clearance of aggregates accumulated in the periplasm. Thus, GlpG may be part of a quality control checkpoint of FimA subunits, ensuring assembly fidelity of type 1 pili. Overall, our study demonstrates that GlpG plays an essential role in UPEC virulence and may represent a viable drug target for the treatment of bacterial infections, such as UTIs, to combat the rise of antimicrobial resistant bacteria.

## Results

### Proteomic analyses of GlpG identified substrates associated with bacterial virulence

Rhomboid proteases were discovered in the early 2000’s, and *E. coli* rhomboid protease, GlpG, quickly became a key model for understanding the structure and mechanism of this family of intramembrane proteases (Ben-Shem et al., 2007; Maegawa et al., 2007; Wang and Ha, 2007). Despite being the first family member to have the structure solved, the interacting partners, physiological substrate(s) and hence, the function of *E. coli* GlpG remained unknown. Our first focus was to identify binding partners/substrates of GlpG using a 3X FLAG tag genomically inserted downstream of the *glpG* gene to enable affinity pulldown and whole cell proteomics in the presence of non-denaturing detergent **(Figure 1A)**. This revealed that GlpG interacted with several proteins with roles in intracellular secretion and protein turnover when assigned to Clusters of Orthologous Groups (COGs) **(Figure 1B)**, in particular with Sec Y and G of the SecYEG translocon, its associated ATP-dependent motor protein SecA as well as its ancillary proteins YidC, SecD and F. Together they drive Sec-mediated translocation of unfolded proteins across the membrane, a tightly regulated process that exports most secretory proteins in bacteria. GlpG may associate with the Sec translocon to readily cleave misfolded or aggregated proteins to prevent the toxic accumulation of aggregates, as part of a role in protein quality control, a function shared by other rhomboid proteases studied in the current literature. We also demonstrate GlpG association with DegQ and DegP **(Figure 1B)**, proteases with central roles in protein quality control that are closely involved in clearing unproductive or harmful protein intermediates, particularly under stress conditions. There is increasing evidence that both DegP and DegQ interact functionally with the SecYEG translocon (Merdanovic et al., 2011). DegP, in particular, has been shown to degrade preproteins that are stalled or improperly inserted into the membrane, suggesting a surveillance role at or near the translocon (Braselmann et al., 2016).

**Figure 1.**
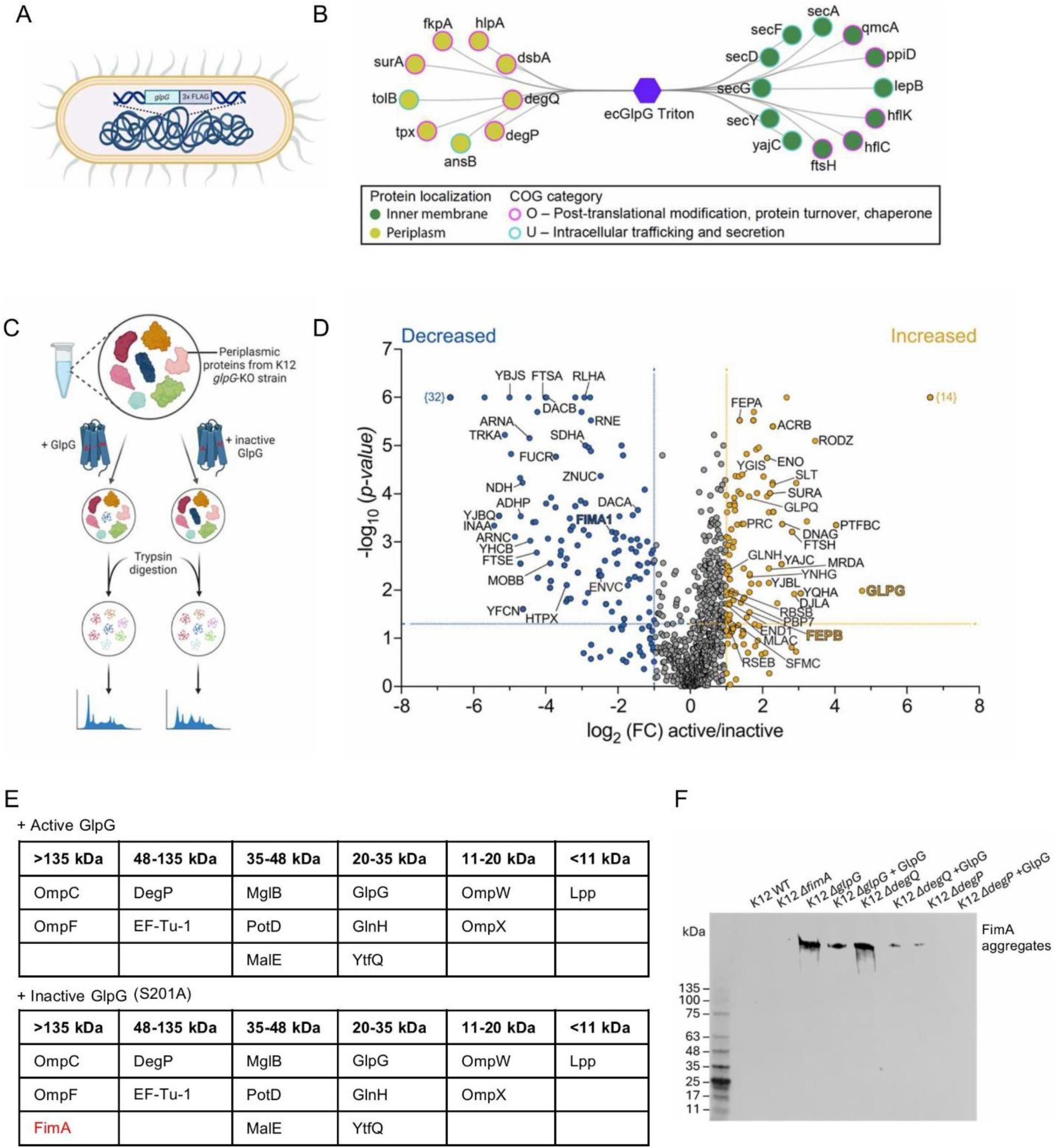
Proteomic analyses of GlpG identified substrates associated with bacterial virulence. A) The *glpG* gene in *E. coli* strain DY330 was genomically tagged with 3X FLAG. B) Cytoscape network of periplasmic and inner membrane proteins identified after affinity pulldown of whole cell extract with *glpG-*3XFLAG followed by LC-MS/MS reveal an enrichment of proteins involved in proteostasis. C) Experimental procedure for LC-MS/MS proteomics of *E. coli* periplasmic proteins. D) The volcano plot showing decreased (blue) and increased (orange) protein abundance levels in extracts after incubation with active relative to inactive GlpG. Lines represent the boundaries for a 2-fold change and a significant p-value of <0.05. Proteins were identified with the UniProt ID. The number of proteins quantified in one sample but not the other is identified in curly brackets. E) In-gel digestion and mass spectrometric characterization of periplasmic proteins of K12 substr. MG1655 *ΔglpG* separated by SDS-PAGE reveal accumulation of aggregated FimA (>135 kDa MW range) when incubated with inactive GlpG, but no accumulation when incubated with active GlpG. F) Western blot of periplasmic proteins separated by SDS-PAGE, detecting FimA aggregates using anti-FimA antibody. Aggregates are defined as >135 kDa. Periplasmic proteins were isolated from K12 wild-type, strains harboring single gene deletions: *ΔglpG, ΔdegQ, ΔdegP*, and rescue strains overexpressing GlpG. K12 *ΔfimA* was used as a negative control.

The list of interacting partners from whole-cell proteomics led us to take a narrower approach to analyze only the resident proteins in the periplasmic space. The active site of GlpG in *E. coli* faces the periplasm, allowing potential substrates to be identified when comparing isolated periplasmic proteins in the presence of active or inactive GlpG using LC-MS/MS **(Figure 1C)**. In addition, this minimizes cytosolic proteins that would otherwise be identified as non-physiological substrates (Quan et al., 2013). Notably, the periplasmic fraction was isolated from a *ΔglpG* mutant of K12 substr. MG1655 *E. coli* to ensure any identifiable proteins were not prematurely cleaved by endogenous GlpG. We hypothesized that proteolysis by GlpG would lead to the generation of smaller protein fragments potentially leading to apparent decrease in protein abundance due to fragments degradation, precipitation in the lysate or running out of the gel. Therefore, proteins identified as significantly decreased in the presence of active GlpG, relative to inactive GlpG, would represent potential cleavage by GlpG **(Figure 1D)**. Several proteins were identified, including those involved in cell division, stress response, and chaperone mediated folding, such as FtsA, RcnB, and PpiD **(Figure S1 and Data S1)**. Importantly, virulence factor FimA, a subunit of type 1 pili, was identified among the proteins significantly decreased in the presence of active protease, drawing our attention in accordance with previous literature, describing colonization defects of pathogenic *E. coli* with a *glpG* knockout strain (Russell et al., 2017).

To further investigate the role of GlpG in proteostasis, an in-gel digestion was performed whereby periplasmic proteins isolated from the K12 *ΔglpG* mutant strain were incubated with active or inactive GlpG, separated by SDS-PAGE and followed by gel excision at different molecular weight ranges to be analyzed using LC-MS/MS. In line with the function of other rhomboid proteases that cleave aggregated substrates to prevent their accumulation (Began et al., 2020; Liu et al., 2020), we sought for a more detailed examination of which proteins may be forming aggregates in the absence of GlpG and could be cleaved by the re-addition of GlpG **(Data S2)**. When periplasmic proteins were incubated with inactive GlpG, FimA was found to be at a higher molecular weight (>135 kDa) instead of its monomeric size (18 kDa) whereas in the presence of active GlpG, this oligomeric form of FimA was not present **(Figures 1E and S2)**. In wild-type (WT) *E. coli* FimA monomers were rarely detected, likely due to their incorporation into pili or rapid turnover **(Figures 1E and 1F).**

In WT *E. coli*, homeostasis of FimA protein in periplasmic space is maintained, as evidenced by the lack of FimA aggregation when performing immunoblot detection by western blot. By contrast, FimA aggregation was detected in the periplasm of K12 MG1655 *ΔglpG*, even under SDS-denaturing conditions **(Figure 1F)**, suggesting that in the absence of GlpG catalytic activity, proteostasis of FimA is dysregulated, causing its accumulation and aggregation within the periplasm during its translocation. When K12 MG1655 *ΔglpG* was transformed with the plasmid bearing active GlpG, we observed notably decreased amount of aggregates, confirming GlpG’s role in protein quality control as seen in many other rhomboid proteases (Began et al., 2020; Liu et al., 2020). To test the hypothesis that GlpG may work in coordination with DegQ and DegP and that these periplasmic proteases may also be involved in the biogenesis of type 1 pili, we detected FimA in the samples of periplasmic proteins isolated from single gene knockout of *degQ* and *degP* in K12 MG1655. Indeed, immunoblot analysis demonstrated aggregated FimA in the periplasm in the absence of either Δ*degQ* and Δ*degP* **(Figure 1F)**, identifying a link between DegQ and DegP proteases and quality control of type1 pili for the first time. The more profound effect of Δ*degQ* on FimA aggregation could be explained by the fact that Δ*degP* protease is not constitutively expressed under normal growth conditions (Kim and Sauer, 2014), as opposed to Δ*degQ*, which has relatively constant level regardless of specific environmental conditions (Farn and Roberts, 2004). Moreover, the K12 MG1655 *ΔdegP* and *ΔdegQ* mutants transformed with plasmid expressing active GlpG exhibited reduced amount of FimA aggregates. This not only suggests that the interplay between GlpG and DegQ/P may reflect a specialized proteostasis module but also places GlpG in a first-responder role to periplasmic aggregation of pilin subunits.

### GlpG activity is essential for pili formation

FimA protein is the structural subunit of type 1 pili, comprising of up to 3000 subunits in a single pilus. It is an abundant and essential component for bacterial adherence to host cells, therefore maintaining its protein homeostasis is crucial in pathogenic *E. coli*. The main function of adhesion virulence factors, such as type 1 pili, is to facilitate attachment of the bacterium to the host cell, often as an initial step in invasion or uptake. To determine if the lack of GlpG catalysis affects type 1 pili formation, transmission electron microscopy (TEM) was used to visualize surface pili. UPEC strain NU149 was chosen to represent a clinically relevant isolate from a patient presenting cystitis (Schaeffer et al., 1987). We show that significantly more surface pili are present on NU149 wild-type relative to a genomically inactive *glpG* strain generated using CRISPR/Cas9, which harbors a Ser201Ala point mutation at the catalytic serine **(Figures 2A**, **2B, S4)**. A genomically inactive *glpG* mutant, hereby denoted as *glpG*-S201A, was used to gain a more physiological insight into the role of the protease compared to a gene knock-out. The growth curve and *glpG* mRNA expression of the resulting strain endogenously expressing catalytically inactive GlpG was unaffected compared to its wild-type counterpart **(Figure S3)**. Importantly, it is known that mutation of the catalytic serine does not disrupt the folding of GlpG protease (Vinothkumar, 2011). The significant impairment of pili formation observed in *glpG*-S201A mutant was rescued when the active GlpG protease was reintroduced to NU149 *glpG*-S201A using an inducible plasmid, with an apparent increase in not only pili but flagella, which was unexpected. The same trends were observed when *fim* protein chaperone, FimC, was overexpressed, assisting the folding, binding and translocation of FimA **(Figures 2A**, **2B and S5)** (discussed in detail below). To test if GlpG expression may be elevated in pathogenic *E. coli* relative to commensal *E. coli,* we quantified *glpG* mRNA transcript levels using RT-qPCR, and indeed detected higher expression in NU149 and J96 UPEC strains compared to K12 MG1655. This result was not affected by different nutrient environments such as LB, high nutrient terrific broth (TB) or M9 minimal media that could influence bacterial growth rate, biomass yield, gene expression, and protein production **(Figure 2C).**

**Figure 2.**
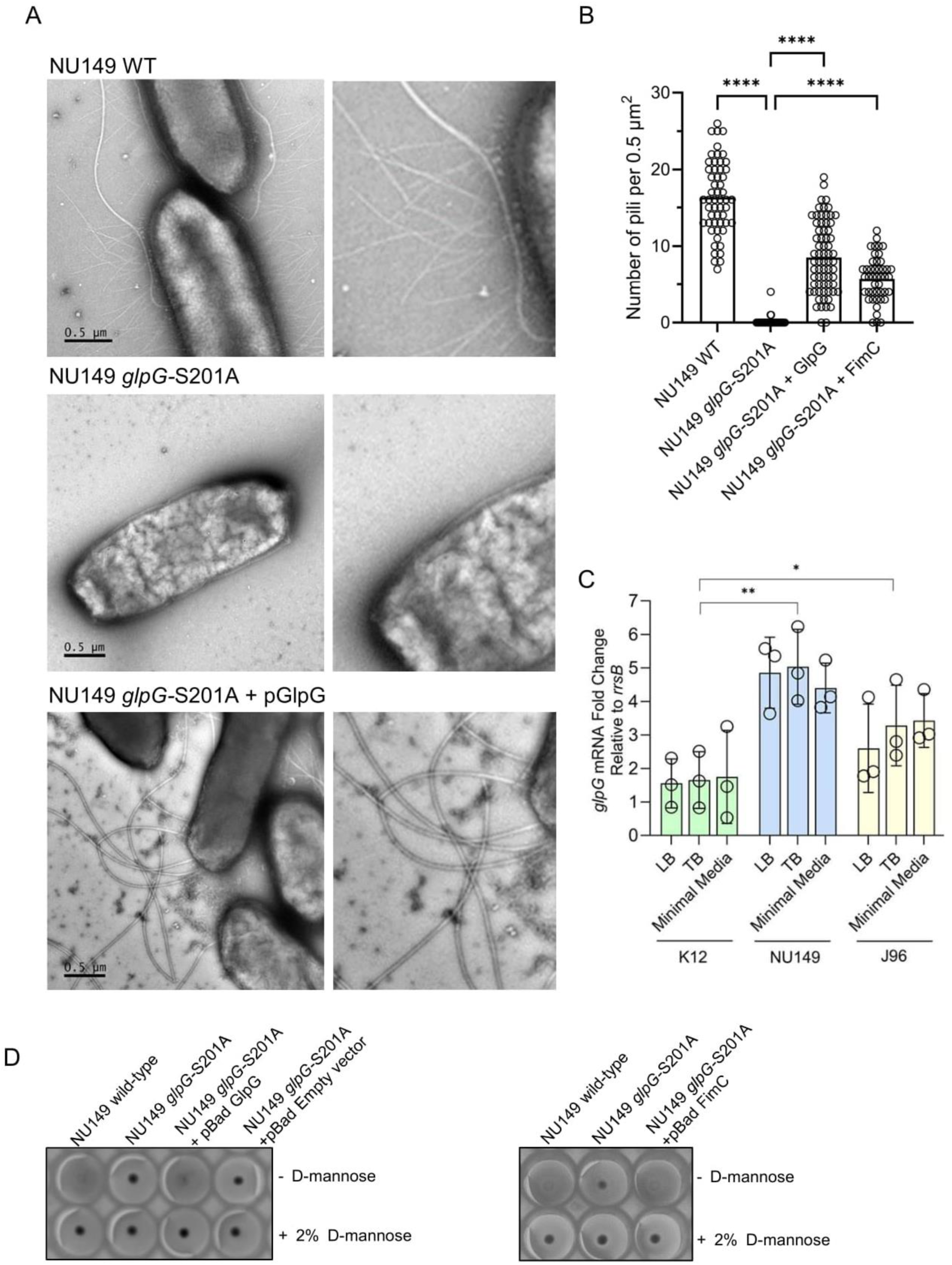
Surface organelle formation is impaired when GlpG activity is not present in UPEC strain NU149. A) Negatively stained electron micrographs of uropathogenic *E. coli* strain NU149 wild-type, the CRISPR/Cas9 *glpG*-S201A mutant, and *glpG*-S201A mutant transformed with inducible plasmid expressing active GlpG. Bar represents 0.5 µm B) Pili from micrographs were quantified by manual counting using n≥30 of *E. coli* cells per strain. Significant differences between strains were calculated using one-way ANOVA. ****P<0.0001 C) Quantification of *glpG* transcription using RT-qPCR analysis in commensal (K12) and uropathogenic (NU149 and J96) *E. coli* strains under different growth conditions. *glpG* mRNA is reported relative to housekeeping gene *rrsB*. Each strain was performed in technical triplicates and biological triplicates. Significant differences were determined using one-way ANOVA. **P<0.01, *P<0.05 D) Hemagglutination assay of NU149 *E. coli* strains confirm *glpG* mutant strains do not possess type 1 pili. The absence of type 1 pili on *E. coli* is confirmed lack by binding to erythrocytes, forming a dense button of erythrocytes at the bottom of the well. As a control, hemagglutination was inhibited by the presence of 2% of D-mannose prior to the addition of bacterial cultures.

To confirm that the surface organelles detected using TEM were indeed type 1 pili, we performed a classical functional test for the presence of type 1 pili: a mannose-sensitive hemagglutination (MSHA) assay on guinea pig erythrocytes. Type 1 pili mediate the hemagglutination (HA) that occurs when piliated bacteria are mixed with erythrocytes of certain species. Since type 1 pili on UPEC bind to mannosylated proteins on host cells, particularly mannose found on uroplakins and integrins by FimH adhesion, the HA reaction can be inhibited by addition of 2% mannose (Evans et al., 1979). The NU149 strains described above were grown under the same conditions as for TEM imaging and assessed for HA **(Figure 2D)**. NU149 wild-type and *glpG*-S201A strain overexpressing active GlpG protease were MSHA positive (type 1 piliated), whereas inactive glpG mutant as well as the cells transformed with an empty pBad vector did not exhibit HA, suggesting the absence of type 1 pili, in agreement with the TEM results. Overall, this data shows that the catalytic activity of GlpG influences type 1 pili, a critical virulence factor in *E. coli*, through the quality control of FimA.

### GlpG activity influences pathogenic *E. coli* invasion

UPEC can infect both bladder and kidneys cells and account for the vast majority of all UTI infections (Terlizzi et al., 2017). To determine whether loss of pili observed in *glpG-*S201A strains results in reduced pathogenicity, an *in vitro* gentamicin protection assay was used to quantify bacterial entry into epithelial cells **(Figure 3A)**. The *glpG*-S201A mutants of UPEC NU149 and commensal K12 substr. MG1655 showed significantly reduced invasion relative to their wild-type counterparts in both T24 human bladder **(Figure 3B)** and mIMCD-3 mouse kidney cell lines **(Figure 3C)**. Notably, this reduction in invasion was rescued to wild-type levels when active GlpG was reintroduced using an inducible plasmid, confirming the loss of GlpG catalysis results in reduced pathogenicity in uropathogenic and commensal strains of *E. coli*.

**Figure 3.**
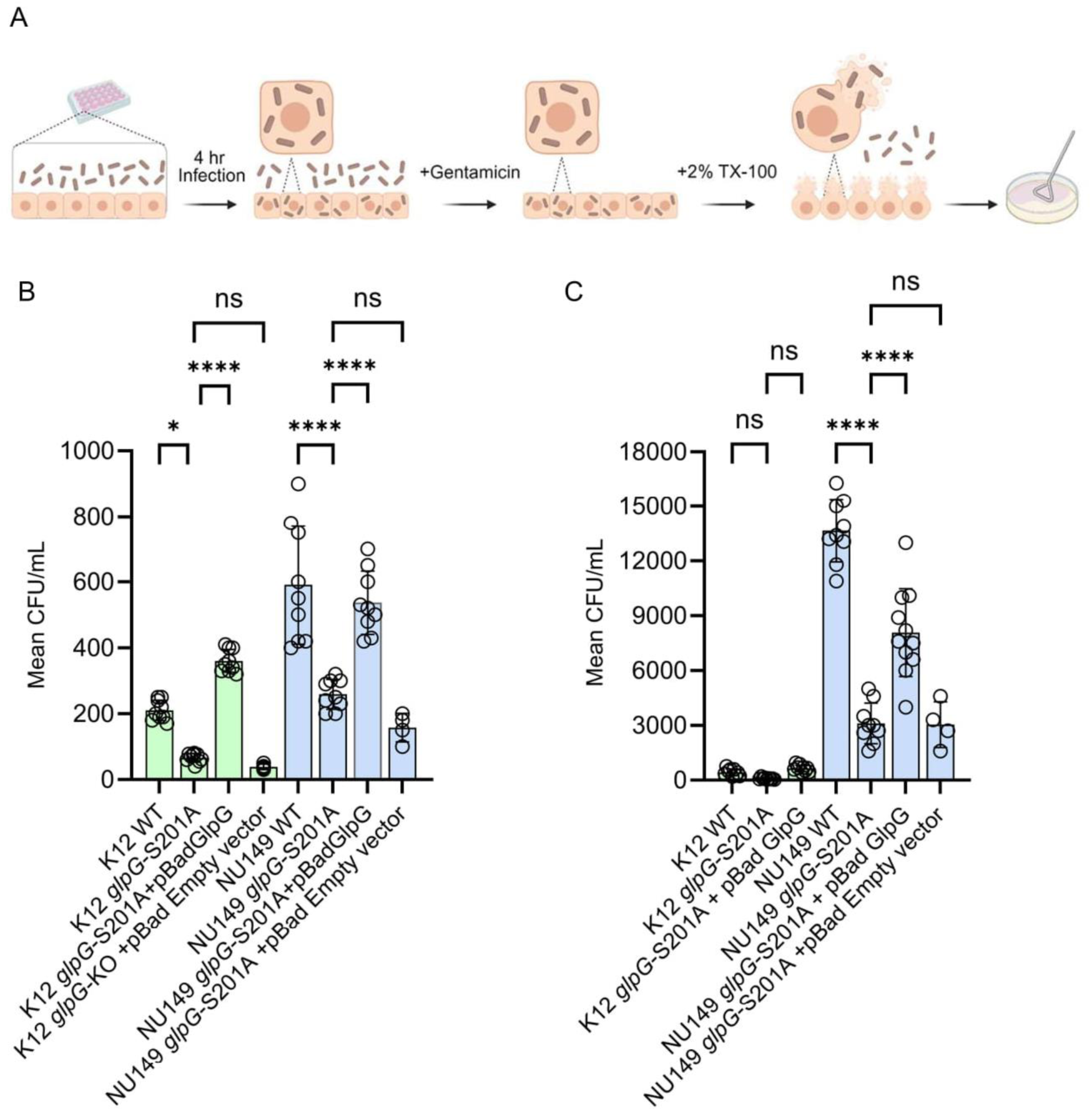
GlpG catalysis is required for bacterial invasion by commensal and uropathogenic strains in bladder and kidney cell lines. A) Gentamicin protection assay used to quantify levels of invasion between different strains of *E. coli* B) Quantification of infection between different strains of *E. coli* against mIMCD-3, mouse kidney cell line and C) T24, human bladder epithelial cell line. Epithelial cells were infected using wild-type commensal K12 substr. MG1655 (green), or UPEC NU149 strains (blue) and their *glpG* mutants. Each strain was performed in technical triplicates and with an n=9. Significant differences between data sets were determined using one-way ANOVA. *:P<0.05, ****:P<0.0001, ns: P>0.05.

Proximal and distal tubular epithelial cells of the kidney have the capability to phagocytose *E. coli* present in the tubular lumen, causing peritubular inflammation (Maesaka and Shimamura, 1984). To determine if the GlpG protease impacts the *E. coli* infectivity for tubular epithelial cells, human kidney organoids differentiated to express segments of the proximal tubule, were infected with NU149 wild-type or NU149 *glpG-*S201A *E. coli*. Kidney organoids infected with wild-type NU149 had *E. coli*-FITC stained structures present within LTL-Cy5 positive regions which label proximal tubules **(Figures 4A**, **4B and S6)**. Meanwhile, a 3.5-fold reduction of detectable bacteria at 469 NB100-FITC per mm^2^ of LTL-Cy5 for NU149 *glpG*-S201A was observed compared to wild-type infected kidney organoids which had 1650 anti-*E. coli*-FITC per mm^2^ of LTL-Cy5 **(Figure 4C)**. Consistent with our results of the gentamicin protection assay, proximal tubular cell invasion for NU149 *glpG-*S201A was significantly lower compared to wild-type, further highlighting the importance of GlpG for bacterial pathogenicity.

**Figure 4.**
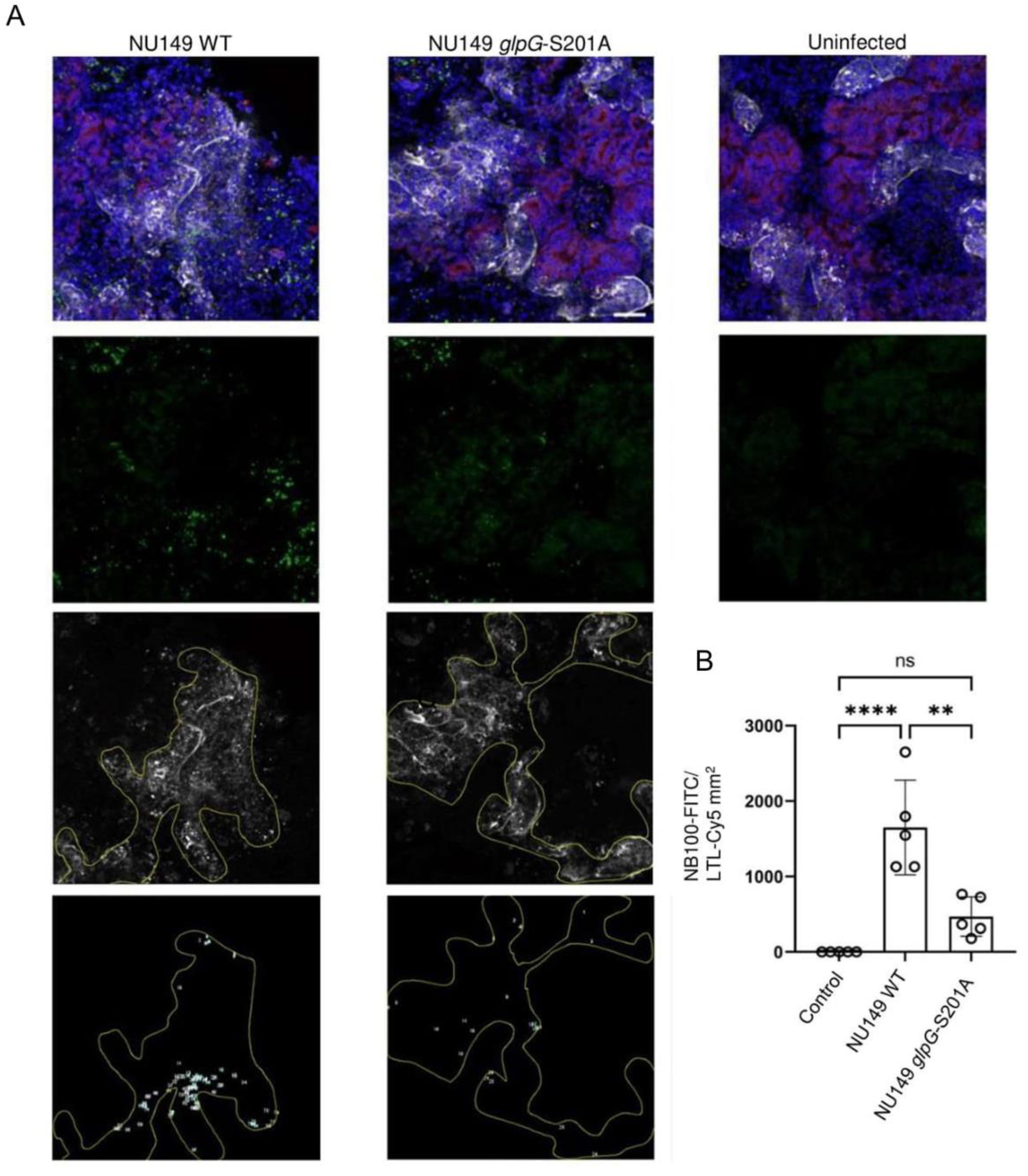
*E. coli* NU149 *glpG*-S201A mutant is unable to invade kidney organoid tubules. A) Immunofluorescence images of kidney organoids (100K) infected with wild-type NU149 or mutant strain NU149 *glpG*-S201A at a ratio of 1:100. *E. coli* were imaged with NC100-FITC antibody detecting O or K antigen serotypes in green, cell nuclei in blue, and proximal tubule visualized with LTL-Cy5. B) Quantification of *E. coli* infection within tubule markers show significantly higher bacterial invasion in the wild-type NU149 sample. Each strain was performed in technical triplicates and with an n=5. Significant differences between data sets were determined using one-way ANOVA. **P<0.01, ****P<0.0001, ns: P>0.05.

### GlpG influences biofilm formation of uropathogenic *E. coli*

Biofilms are the main cause of recurrent bacterial infections, largely because their environment protects the pathogen from administered antibiotics (Flemming et al., 2016). Previous work showed that pathogenic *E. coli* can form biofilms on the surface of epithelial cells, which promote their persistence *in vivo* (Elhenawy et al., 2021). Importantly, surface pili were shown to play a key role in driving the formation of *E. coli* biofilms (Elhenawy et al., 2021; Ghigo, 2001; Patkowski et al., 2023). Given its role in the homeostasis of type 1 pili, we reasoned that GlpG is required for the assembly of biofilms by UPEC on biotic surfaces. To test this, we infected mIMCD-3 kidney cells with NU149 wild-type or *glpG*-S201A isogenic strains for 6 hours to allow biofilm formation. To quantify these structures, we tallied the number of bacterial aggregates larger than 100 µm^2^, which was previously shown to correspond to the sizes of biofilms made by pathogenic *E. coli* (Elhenawy et al., 2021). Interestingly, we observed a significant reduction in biofilm formation (∼25-fold) by the mutant relative to the wild-type **(Figure 5)**. This was rescued by overexpressing GlpG in the mutant strain, confirming the role of this protease in facilitating biofilm assembly, likely through its role in FimA proteostasis.

**Figure 5.**
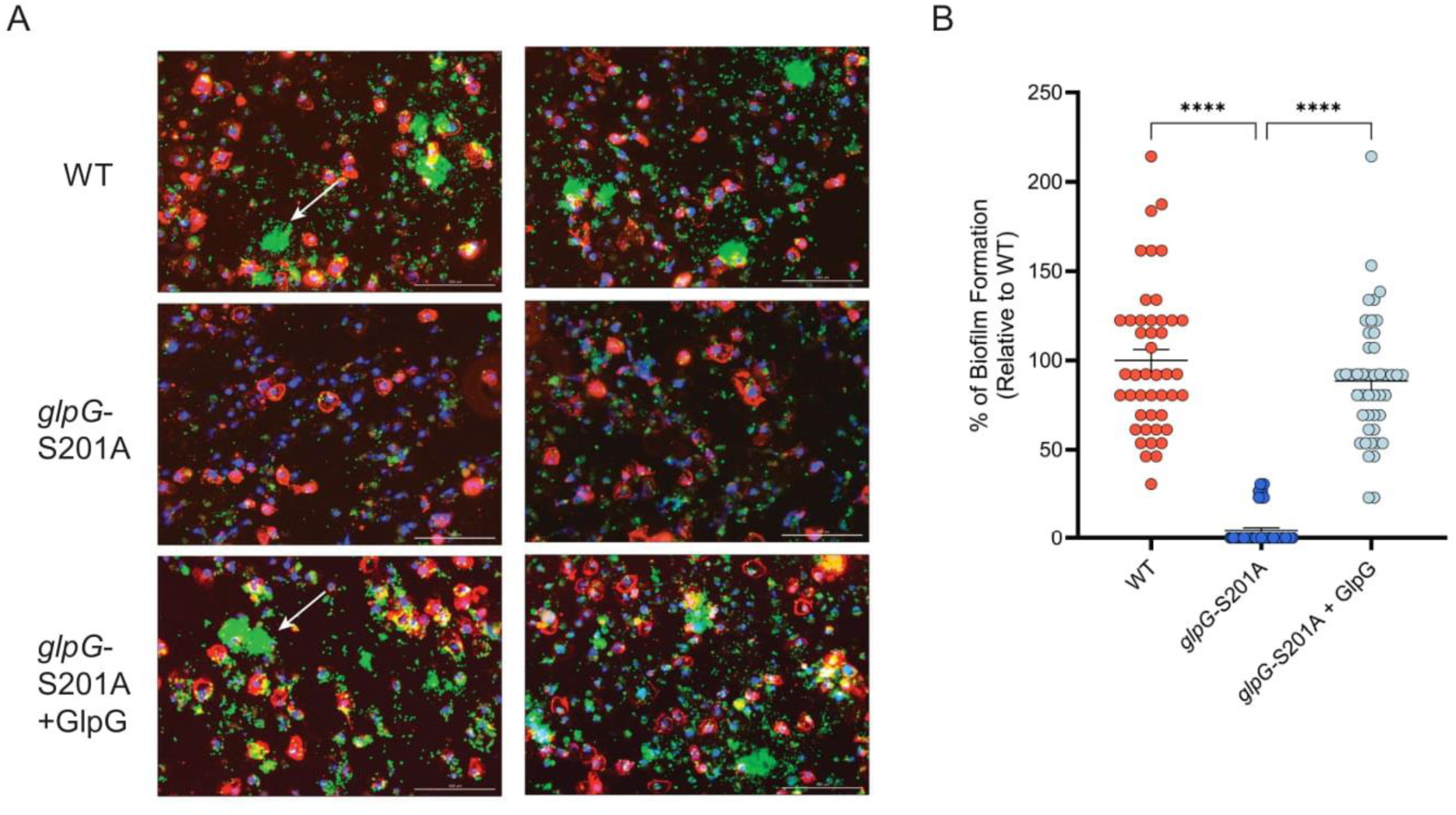
GlpG is required for biofilm formation by uropathogenic *E. coli*. A) Immunofluorescence staining shows mIMCD-3 epithelial cells in red with DAPI stained nuclei in blue, infected with different UPEC stains in green. White arrows indicate examples of biofilms. Each panel is representative of 45 images per group obtained from 3 biological replicates. Scale bar is 100 µm B) Quantification of biofilm formation by the different UPEC stains relative to wild-type. ****P<0.0001, Kruskal-Wallis test.

Another method to validate the effect GlpG has on the formation of bacterial biofilm is safranin staining, used to quantify biofilm mass produced in UPEC on an abiotic surface. Indeed, the NU149 *glpG*-S201A mutant, which displays impaired pili formation, showed significant reduction in biofilm production compared to the wild-type **(Figure 6A)**. Interestingly, the *glpG*-S201A strain overexpressing active GlpG protease showed a significant increase in biofilm biomass. These results demonstrate a correlative relationship between the catalytic activity of GlpG, pili formation and biofilm forming abilities of *E. coli*. Indeed, negatively stained electron micrographs of NU149 demonstrated increased amount of GlpG protease in the cell significantly affected the amount of pili on the surface.

**Figure 6.**
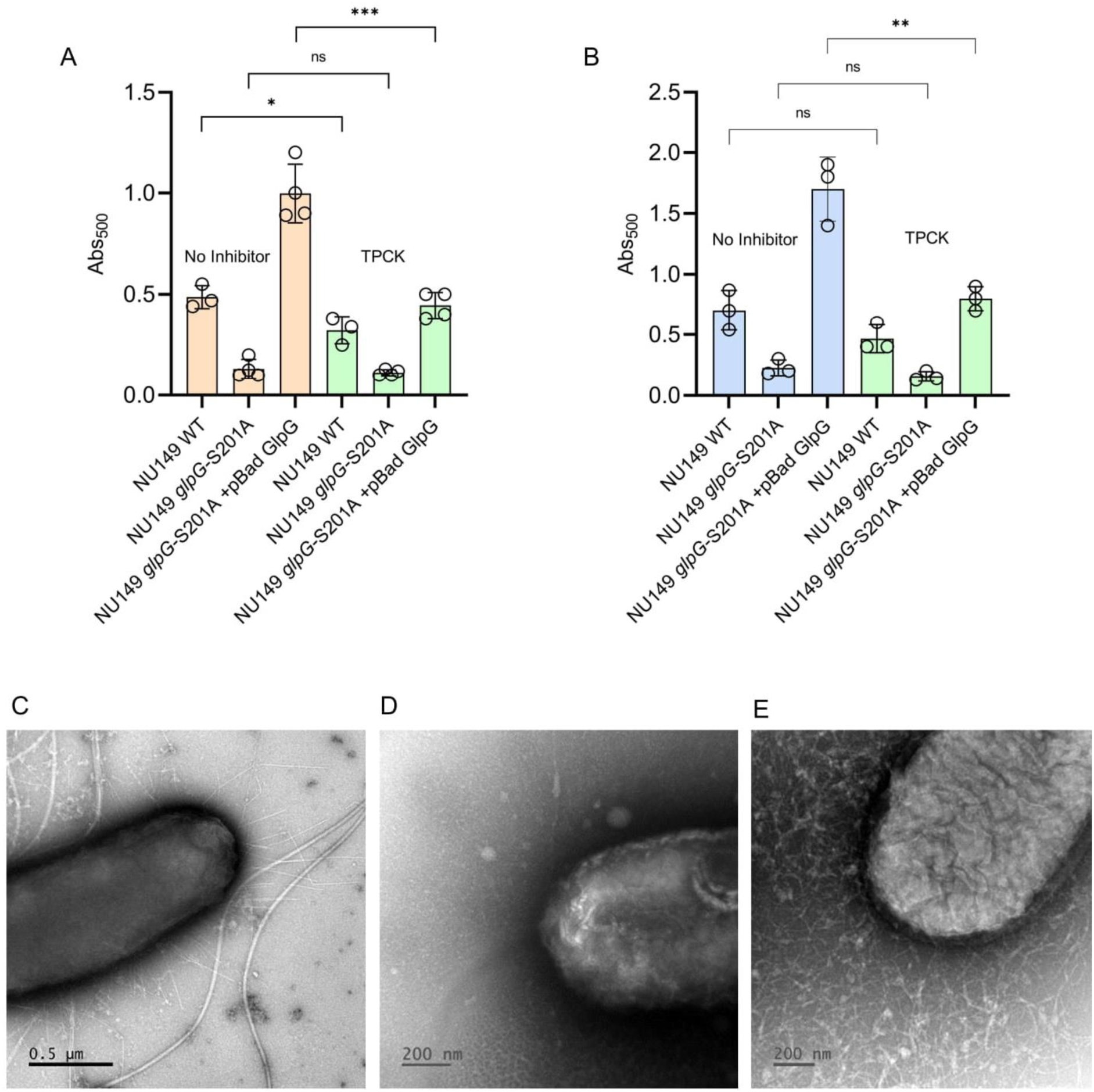
Microtiter biofilm assay of different strains of UPEC NU149. A) Quantification of safranin-stained biofilms between different strains of NU149 with and without treatment with TPCK, the same time as GlpG induction or 2 hours after GlpG induction (B), measured by absorbance at 500 nm. The data represents the mean ± SEM of three independent experiments performed in duplicate. Significant differences between data sets were determined using one-way ANOVA. *P<0.05, **P<0.01, ***P<0.001, ns: p>0.05 C) Electron micrographs of negatively stained NU149 *glpG*-S201A +GlpG cells without inhibitor and when TPCK was added the same time as GlpG induction (D) or 2 hours after GlpG induction (E). Each strain was performed in technical triplicates and biological triplicates.

### GlpG inhibition affects the expression of surface virulence factors

To investigate whether the expression of surface virulence factors can be blocked by inhibiting *E. coli* GlpG rhomboid protease, we used commercially available serine protease inhibitor, tosyl phenylalanyl chloromethylketone (TPCK). We determined the inhibitory concentration of TPCK for GlpG *in vitro* using a peptidic FRET substrate (Mca-FATA*AFGSP-Dpn) designed by our team based on the cleavage sequence of the transmembrane domain of TatA protein from *P. stuartii* – a known bacterial substrate of rhomboid protease AarA that *E. coli* GlpG is also able to cleave (Strisovsky et al., 2009). We measured the IC_50_ value of TPCK for GlpG to be 296 µM **(Figure S7)**, consistent with previous observations (Zoll et al., 2014).

To explore the effect of TPCK on virulence factor expression, bacterial adherence and formation of biofilm, biofilm inhibition and biofilm eradication assays were performed. In parallel, electron micrographs of *E. coli* cells treated with TPCK were taken to observe changes in surface organelles presentation. Biofilm inhibition was assessed by adding the inhibitor at the same time as the re-addition of active GlpG to the *glpG*-S201A strain (NU149 *glpG*-S201A +GlpG) and assessing the biofilm production. Overall, we observed a decrease in biofilm formation in the presence of TPCK for NU149 wild-type, but a more profound inhibitory effect was observed in NU149 *glpG*-S201A +GlpG, which we show produces more biofilm than the wild-type **(Figures 6A and 6D)**. We then explored whether inhibition of GlpG can aid in biofilm eradication, or the removal of pili that have already formed. To test this, biofilms were first pre-established with bacterial growth before the addition of TPCK inhibitor. When quantified, we did not observe a significant difference in adhered biomass for wild-type NU149 treated with TPCK, however the NU149 *glpG*-S201A +GlpG strain, again, demonstrated a significant decrease in biofilm mass in the presence of TPCK **(Figures 6B and 6E)**. To ensure the reduction in biofilm was not due to toxic effects of TPCK, a metabolic activity assay was performed in parallel. Using tetrazolium salt dye, 2,3,5-triphenyl tetrazolium chloride (TTC), detecting cell redox activity and viability, we did not observe the decrease in metabolism for any of the strains used, both with and without inhibitor treatment **(Figure S8)**, indicating that biofilm eradication was not in fact due to cell death.

These observations were confirmed with negatively stained electron micrographs of NU149 *glpG*-S201A +GlpG. When TPCK was added along with the inducing agent for GlpG expression, the cells had a drastic decrease in pili (**Figures 6D**), whereas when TPCK was added after surface elements were already formed, we observed a fragmented pili network (**Figure 6E).** Pili are known to be fragile and constantly turned over and if assembly is faulty for reasons such as GlpG inhibition, the resulting pilus rods may have structural defects and reduced stability, making them more prone to dissociation. This indicates that GlpG is important for maintaining the integrity of the pili in pathogenic bacteria, a crucial feature for retaining the biofilm.

### GlpG cleaves aggregated FimA to prevent accumulation in the periplasm

To confirm the mechanism of GlpG and its contribution to the quality control of FimA, we performed an *in vitro* endpoint activity assay capturing GlpG-mediated cleavage of FimA aggregates. Recombinantly expressed active GlpG, either detergent (DDM) solubilized or in proteoliposomes (PLS) reconstituted in *E. coli* lipids to mimic the physiological membrane, was added to the purified periplasmic fraction of the K12 MG1655 *ΔglpG* mutant containing aggregated FimA. The time course cleavage was visualized by immunoblotting with FimA antibody **(Figures 7A and 7B)**. Indeed, upon incubation with active GlpG protease in DDM and in PLS, an observed decrease in FimA aggregates was evident over time confirmed by densitometry analysis of the corresponding bands **(Figures 7C and 7D),** demonstrating GlpG can cleave FimA aggregates in the lipid bilayer.

**Figure 7.**
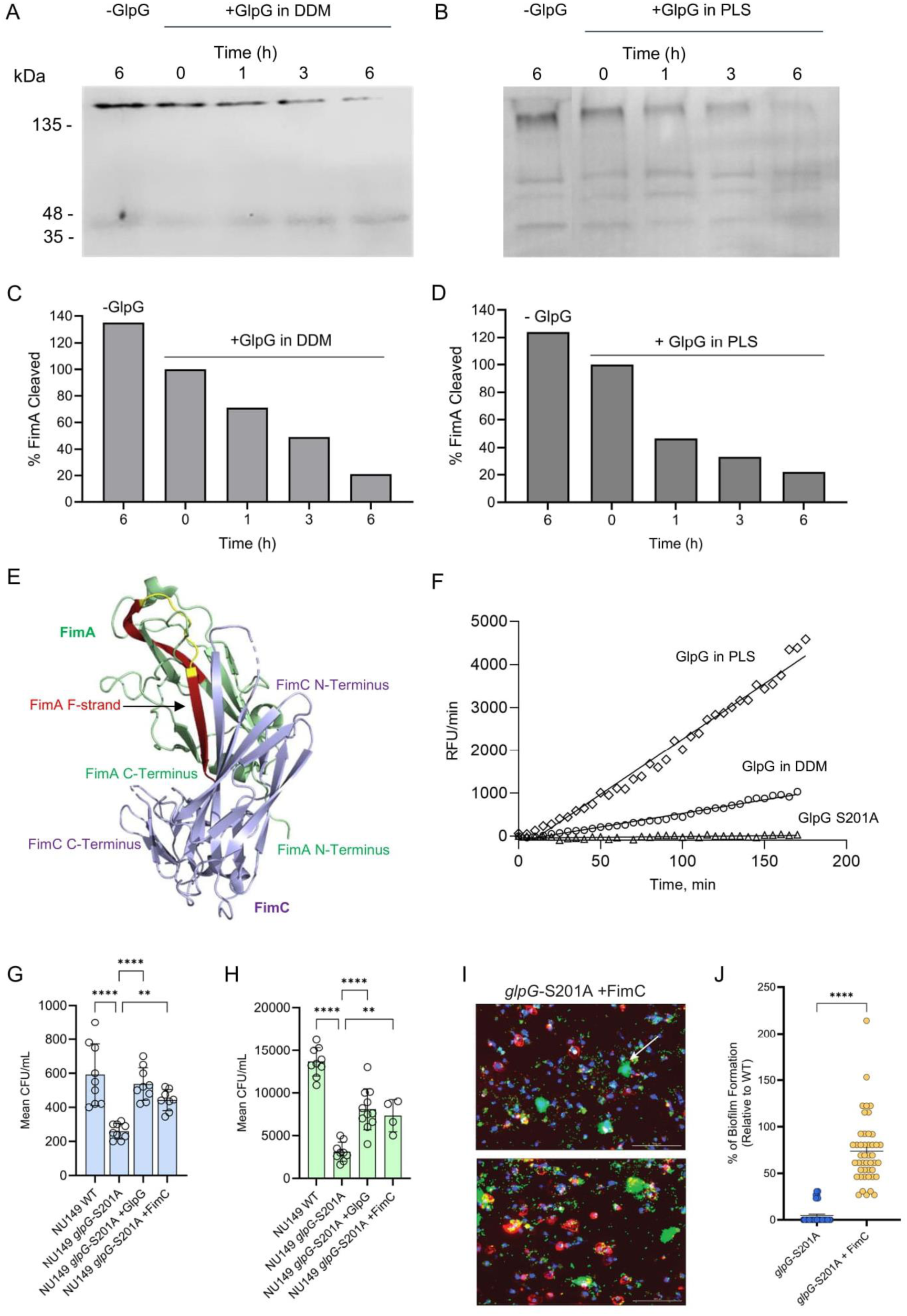
GlpG cleaves FimA at the C-terminal tail for subsequent degradation to prevent accumulation of aggregates in the periplasm. A) Western blot analysis of GlpG-mediated cleavage of FimA aggregates. Periplasmic proteins were isolated from K12 substr. MG1655 *glpG*-KO mutant and incubated with purified recombinant GlpG at 37 °C for 6 hr. Aliquots were taken at time points and the cleavage reaction was quenched with SDS-PAGE buffer. The sample of periplasmic proteins incubated without GlpG at 37 °C for 6 hr was used as negative control, evidenced by the no observed self-disruption of aggregates. FimA antibody was used for visualization. Quantification by densitometry analysis of bands of aggregated FimA (B) and monomeric form (C) over time. D) Time course of GlpG-mediated cleavage of FRET-FimA peptide. 0.5 µM of GlpG was incubated with 50 µM of peptide at 37 °C; reaction was monitored by measuring fluorescence at 490 nm. E) The structure of FimA (green) complexed with FimC chaperone (purple) (PDB code 4DWH; Crespo et al., 2013)), showing the interaction of the C-terminal end of FimA with the FimC chaperone. The C-terminal peptide is shown in red with putative GlpG cleavage sites colored in yellow. F) Quantification of infection against mIMCD-3 cells of NU149 *glpG*-S201A overexpressing FimC compared to wild-type, NU149 *glpG*-S201A, NU149 *glpG*-S201A overexpressing GlpG. Each strain was performed in technical triplicates and with an n=9. Significant differences between data sets were determined using one-way ANOVA. ****:P<0.0001, ****:P<0.0001, ****:P<0.0001 G) Immunofluorescence staining shows mIMCD-3 epithelial cells in red with DAPI stained nuclei in blue, infected with NU149 *glpG*-S201A + FimC. Panel is representative of 45 images per group obtained from 3 biological replicates. Scale bar is 100 µm B) Quantification of biofilm formation by the different UPEC stains relative to wild-type. ****P<0.0001, Kruskal-Wallis test.

During FimA translocation through the Sec pathway, FimA has an exposed, hydrophobic C-terminal tail that is prone to aggregation in an aqueous periplasm. This C-terminal tail of FimA must immediately bind to FimC chaperone in the periplasm for its eventual transport to the outer membrane. Thus, if after SecYEG-export to periplasm FimA is improperly folded, or when insufficient FimC chaperone molecules are available to bind and transport FimA through the periplasm, pilins will misfold and aggregate and are targeted by DegP and DegQ, the housekeeping periplasmic proteases. GlpG most likely influences this pathway by processing or regulating misfolded pilins upstream of DegP/Q-mediated degradation GlpG likely maintains proteostasis by cleaving the C-terminus of FimA subunits and “untangling” big oligomers. To verify the cleavage site in the FimA molecule, we designed a FRET-FimA peptidic substrate (DABCYL-QARYFATGAATPGAANADATFKVQYQ-Glu(EDANS) based on the sequence of the FimA peptide identified in LC-MS/MS analysis of periplasmic proteins. The identified sequence is located on the C-terminus of FimA, residues 157 to 182, which forms the hydrophobic groove and binding site for FimC chaperone during translocation **(Figure 7E)**. Due to the hydrophobic nature of the region, the FRET-FimA peptide was prone to aggregation and had limited solubility, as expected. Cleavage of the peptide was demonstrated using purified GlpG in DDM and PLS with no cleavage by catalytically inactive GlpG as a control **(Figure 7F)**.

To confirm the role of GlpG in maintaining the fine balance between free FimA that is prone to aggregation, and FimA bound to FimC for pilin assembly, we investigated the effect of increasing the ratio of FimC relative to FimA in the NU149 *glpG*-S201A strain on cell invasion and biofilm formation. NU149 *glpG*-S201A, which we show has reduced pili and invasion of host cells, had pili **(Figure 2)**, as well as host invasion **(Figures 7G and 7H)**, return to wild-type levels when overexpressing FimC and. This is further supported by the partial rescue of biofilm formation upon overexpressing FimC in the *glpG*-S201A mutant where we observed ∼75% restoration of biofilm biogenesis, suggesting that overexpression of this chaperone can prevent the aggregation of pilin subunits when GlpG is absent **(Figures 7I and 7J)**. This change in equilibrium, increasing available FimC to bind free FimA, rescues the pili-expressing, invasive phenotype. Since FimA requires FimC to reach the outer membrane, having excess chaperone available to bind and transport FimA during translocation limits the presence of unbound FimA and potential aggregation in the absence of GlpG proteostasis **(Figure 8)**. By contrast, when the cell has less FimC relative to FimA during type 1 pili formation, GlpG performs its role in protein homeostasis, cleaving FimA at the exposed, hydrophobic C-terminal tail. By identifying FimA as a physiological substrate of *E. coli* GlpG, our data supports the role of GlpG in maintaining type 1 pili quality control by cleaving FimA aggregates for subsequent degradation by other proteases such as DegP and DegQ that were identified as interactors of GlpG in whole cell proteomics. This prevents the accumulation of FimA in the periplasm and obstruction of the translocation pathway during type 1 pili assembly.

**Figure 8.**
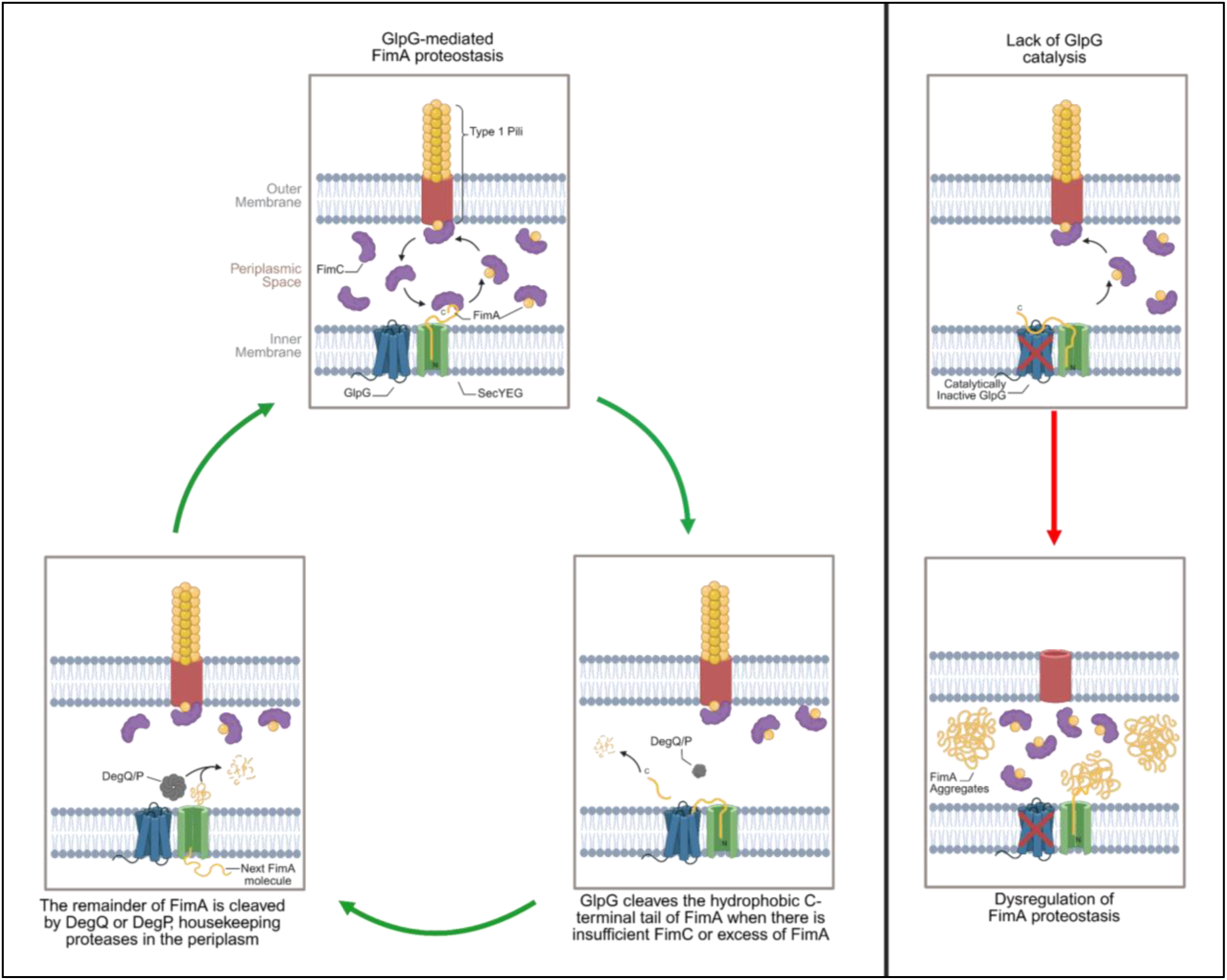
Summary of the role of GlpG in maintaining protein quality control of FimA for the proper assembly of type 1 pili in *E. coli*. GlpG (blue) is closely associated with members of the SecYEG translocon (green) as well as other resident proteases in the periplasm such as DegQ and DegP (grey) to readily prevent and clear FimA (yellow) aggregates. When FimC chaperone (purple) is not available to bind FimA, GlpG cleaves FimA at its hydrophobic C-terminal tail to facilitate its clearance and prevent the aggregation of FimA in the periplasm. The remainder of the FimA molecule is degraded by known housekeeping proteases that cleave misfolded proteins in the periplasm, DegP and DegQ. Loss of GlpG catalytic activity causes the accumulation of FimA aggregates and impaired type 1 pili formation, which in turn affects the pathogenicity of *E. coli*.

## Discussion

Our study provides mechanistic insight into a previously observed phenotype where deletion of the *glpG* gene significantly impaired ExPEC persistence in the murine intestine (Russell et al., 2017). ExPEC use a variety of structural (pili, curli and flagella) and secreted (toxins and iron-acquisition systems) virulence factors to disseminate and persist in sterile niches such as the urinary tract, bladder and kidney. In type 1 pili, a single pilus contains thousands of repeating FimA monomers, which assemble non-covalently, extending outwards from the cell. Being a hydrophobic protein and physiologically high in abundance necessitates tightly regulated translocation and proteostasis mechanisms to ensure that any misfolded FimA is cleared from the cell.

Rhomboid proteases are known to cleave transmembrane substrates (Liu et al., 2020; Maegawa et al., 2007). In this study, we identified FimA as the first physiological substrate for *E. coli* GlpG, shifting the paradigm for intramembrane proteases beyond sole recognition of transmembrane substrates and opening the door to explore more novel soluble substrates for this superfamily of proteases. Moreover, we identified several proteases, DegP and DegQ, that work in concert with GlpG and are also involved in protein quality control in *E. coli* periplasmic space by removing misfolded and aggregated proteins (Kolmar et al., 1996), though it is unclear if DegQ or DegP physically interact with GlpG.

We showed that GlpG cleavage site is located on the C-terminus of the FimA molecule, a region critical for proteostasis of FimA subunits. As individual unfolded FimA is transported across the inner membrane via the SecYEG translocon, it is immediately bound by chaperone FimC. Upon binding, the chaperone donates its β-strand to interact with a hydrophobic groove located between N- and C-terminal β-strands within the FimA molecule. This event accelerates the folding of FimA and prevents premature assembly of the subunits in periplasm (Vetsch et al., 2006). The same hydrophobic groove interacts with the incoming FimA subunit during pili assembly on the outer membrane, where the donor β-strand of FimC is replaced by β-strand of the next FimA molecule (Puorger et al., 2011). However, if upon FimA-FimC association the folding of FimA is unsuccessful or if two orphan FimA molecules form a complex prematurely in periplasmic space, it can lead to accumulation of non-functional proteins. We propose that the cleavage of the C-terminal peptide in FimA molecule by GlpG disrupts the binding site for both FimC and the donor FimA, making FimA accessible for further degradation. Thus, GlpG cleaves membrane-proximal misfolded FimA clusters, reducing aggregation or converting them into smaller fragments. These fragments or unassembled pilins are then recognized and degraded by DegP/DegQ in the periplasm. Our data supports physiological co-localization of GlpG and these proteases required for protein trafficking. Together, they form a proteostasis network that surveils and clears misassembled FimA, preventing toxic accumulation and allowing pilus biogenesis to proceed correctly. Indeed, this is in line with previous research on eukaryotic rhomboid protease Der1 and its role as a key regulator in endoplasmic reticulum (ER) associated protein degradation (ERAD) as a member of a ubiquitin ligase complex that degrades misfolded proteins in the ER upon its retrotranslocation into the cytosol (Wu et al., 2020). Interestingly, this is the first evidence of DegP and DegQ involvement in type 1 pili proteostasis. DegP has been identified as the protease that maintains the homeostasis of P pilus by cleaving pilin, PapA subunits, whereas *degP* mutant strains were found to accumulate higher levels of pilin subunits in the periplasm (Hal Jones et al., 1997; Jones et al., 2002).

Type 1 pili was observed to be affected by the absence of GlpG activity, resulting in reduced bacterial invasion and biofilm production. Type 1 pili is known to affect bacterial motility and locating surface contacts, a crucial step in the early stages of biofilm formation. Biofilms are major contributors to bacterial persistence as they provide bacterial communities protection from environmental stress and antibiotics, allowing increased potential for recurrence. Previous studies have also demonstrated that rhomboid proteases affect biofilm formation in *Mycobacterium smegmatis* and *H. volcanii*, showing less biofilm produced when one of the rhomboid genes was deleted (Costa et al., 2025; Kateete et al., 2012). Similarly, we show reduced biofilm formation on biotic and abiotic surfaces when *glpG* is inactivated in *E. coli*, in agreement with the role of GlpG in maintaining pilin homeostasis. Interestingly, biofilm formation was significantly increased when GlpG protease was expressed beyond wild-type levels. Importantly, overexpressing FimC rescued biofilm formation in the absence of a functional GlpG. This result further supports our model in which GlpG plays a key role in eliminating misfolded pilin subunits.

As the threat of antimicrobial resistant pathogens becomes increasingly more prevalent, surpassing multiple classes of antibiotics, new strategies that can help treat such bacterial infections are crucial. The gene encoding bacterial rhomboid protease, *glpG*, is evolutionarily widespread among prokaryotes and structurally conserved within the active site housing the serine-histidine catalytic dyad embedded in the plane of the membrane. Our results clearly show rhomboid protease activity is essential for generating virulence factor type 1 pili in UPEC and that the expression of type 1 pili can be blocked by inhibition of the GlpG protease. Currently new antibacterials are in preclinical and clinical trials, many of which target bacterial proteins to disrupt the function of virulence factors. GlpG therefore represents a strong target in the development of antivirulents to combat AMR given its critical role in type 1 pili.

## Supplemental information

Document S1. Figures S1–S9

- DATA AVAILABILITY

- Lead Contact
- Material availability
- Data and code availability
- METHOD DETAILS

- Periplasmic protein isolation
- GlpG Purification
- Reconstitution of GlpG in proteoliposomes
- LC-MS/MS proteomics on periplasmic proteins
- LC-MS/MS in-gel digestion proteomics on periplasmic proteins
- FimA cleavage time point assay
- Western blot
- *In vitro* cleavage assay
- Construction of mutant strains
- Negatively stained transmission electron microscopy of *E. coli*
- Hemagglutination assay
- RT-qPCR of *glpG*
- Mammalian cell culture conditions
- Gentamicin protection assay
- Infecting kidney organoids with *E. coli*
- Immunofluorescence and confocal microscopy of kidney organoids
- Quantification of bacteria infected proximal tubules
- GlpG inhibition assay
- Biofilm formation assay on biotic surfaces
- Biofilm formation assay on abiotic surfaces
- Biofilm inhibition assay
- Biofilm eradication assay
- Metabolic activity assay
- LC-MS/MS proteomics on whole cells

## DATA AVAILABILITY

### Lead Contact

Further information and requests for reagents may be directed to the lead contact Joanne Lemieux.

### Materials availability

Reagents and strains generated in this study are available, through request to the lead contact with a completed Material Transfer Agreement.

### Data and code availability

- Proteomic data are available as supplemental information with this manuscript.
- This paper does not report original code.
- Any additional information required to reanalyze the data reported in this paper is available from the lead contact upon request.

## METHOD DETAILS

### Periplasmic protein isolation

Periplasmic proteins were isolated using methods described by previous groups (Quan et al., 2013). K12 substr. MG1655 *ΔglpG* mutant was grown to OD_600_ 1.0 and centrifuged at 5,000 x *g* for 10 minutes at 4 °C. Cells were resuspended in a Tris-Sucrose-EDTA buffer (200 mM Tris pH 8.0, 500 mM sucrose, 1 mM EDTA, 1 mM PMSF) with a cOmplete EDTA-free protease inhibitor cocktail tablet (Roche, Cat. # 11836170001) and incubated on ice for 30 min. The sample was centrifuged 20,000 x *g* for 30 min at 4 °C where the final supernatant contains the periplasmic proteins.

### GlpG Purification

Recombinantly expressed *E. coli* GlpG and inactive mutant GlpG Ser201Ala were purified as previously described by our group (Arutyunova et al., 2017a). Briefly, *E. coli* GlpG-TEV-6XHis or GlpG-S201A-TEV-6XHis in a pBAD vector was expressed in TOP10 cells with 0.002% arabinose at OD_600_ 0.8 and collected by centrifugation at 5,000 x *g* for 15 min at 4 °C. The pellet was resuspended in lysis buffer containing 50 mM Tris, pH 8.0, 200 mM NaCl, 1 mM phenylmethylsulfonyl fluoride, protease inhibitor cocktail tablet (Roche Cat. #), and lysed using high-pressure with an Emulsiflex. The proteins were separated from cellular debris at 13,000 x *g* for 30 min at 4 °C and the resulting supernatant was centrifuged at 40,000 x *g* for 2 hr at 4 °C to isolate the membrane fraction. Membranes were resolubilized in a lysis buffer containing 1% Dodecyl-β-D-maltoside, incubated with Ni-NTA resin and eluted with increasing concentrations of imidazole in the lysis buffer. Fractions containing GlpG-TEV-6XHis were further incubated with TEV protease and incubated in Ni-NTA resin again. The flow-through contains full-length GlpG while TEV-6XHis remains bound to the resin. The resulting protein fraction was concentrated with a 30 MWCO Amicon spin concentrator (Millipore, Cat. #UFC9030) and injected into a HiLoad 16/600 Superdex 200 pg column (Cytiva, Cat. #28-9893-33).

### Reconstitution of GlpG in proteoliposomes

400 μg *E. coli* polar lipids (Avanti) suspended in chloroform were dried under nitrogen stream in a glass tube, followed by overnight incubation in a desiccator to yield a thin film of lipid. 50 μl of water and DDM detergent was added to the lipid film for resuspension at room temperature for 10 min, followed by the addition of purified GlpG protease (400 μg) in 50 mM Tris-HCl, pH 8.0, 150 mM NaCl, 20% glycerol, 0.1% DDM, to yield the final weight ratio of 1:1:2, GlpG:lipid:detergent. The detergent was slowly removed by the addition of SM2 Biobeads (Bio-Rad) while stirring on ice for 6 h to allow for the generation of proteoliposomes (PL); the process was controlled by the regimen of Biobead addition. 50%:20% sucrose density gradient ultracentrifugation was used to purify PL. The concentration of reconstituted GlpG was assessed by BCA assay (Pierce BCA Protein Assay Kit, ThermoFisher) and the purity was confirmed with Coomassie staining after SDS-PAGE.

### LC-MS/MS proteomics on periplasmic proteins

Periplasmic proteins isolated from K12 substr. MG1655 *ΔglpG* mutant were incubated with 5 µM of active or inactive GlpG and separated based on molecular weight using 4-20% Mini-PROTEAN TGX Precast Gels (Bio-Rad, Cat. # 4561094) at either 1) 100 V for 5 min, or a migration of 1.5 cm or 2) 150 V for 30 minutes where proteins were analyzed separately based on molecular weight range. Gels were fixed for 20 min (50% ethanol, 2% phosphoric acid), washed for 20 min in MilliQ H_2_O and stained overnight with a blue silver Coomasie stain (10% phosphoric acid, 20% ethanol, 750 mM ammonium sulfate, 13 mM Coomassie blue G-250) and washed for 20 minutes in MilliQ H_2_O.

The band or bands were excised from the gel and further cut into 1 mm pieces. The gel pieces were transferred to a round bottom 96 well plate for destaining (50 mM ammonium bicarbonate, 50% acetonitrile). The plate was incubated for 10 min at 37 °C. The solution was removed from the wells and the destaining procedure was repeated 4 times. The solution was removed and replaced by acetonitrile for dehydration. The samples were incubated again at 37 °C for 10 min. Dehydration was repeated until the gel pieces became white (3 times). The samples were dried at 37 °C for 10 min, rehydrated in a reducing solution (100 mM ammonium bicarbonate and 10 mM β-mercaptoethanol) and incubated for 30 min at 37 °C. The reducing solution was removed. The samples were alkylated with iodoacetamide (10 mg/mL of iodoacetamide and 100 mM ammonium bicarbonate) and incubated for 30 min at 37 °C. The gel bands were subsequently washed with 100 mM ammonium bicarbonate and incubated for 10 min at 37 °C (repeated twice). The samples were then dehydrated in acetonitrile for 10 min at 37 °C (repeated twice). Acetonitrile was removed and the samples were dried at 37 °C for 10 min. The gel bands were then trypsinized (0.3 μg of mass-spec grade trypsin per well, Promega) overnight at room temperature. The solutions containing tryptic peptides were transferred to a V-bottom 96-well plate. Tryptic peptides were further extracted from the gel with 2% acetonitrile and 1% formic acid followed by incubation at 37 °C for 1 h. The extraction was repeated using 50% acetonitrile and 0.5% formic acid followed by incubation at 37 °C for 1 hr. The samples were dried with vacuum and resuspended in 20 mL of 0.1% formic acid for analysis on liquid chromatography tandem mass spectrometry (LC-MS/MS). Peptides were separated using a nanoflow-HPLC (Thermo Scientific EASY-nLC 1200 or 1000 System) coupled to a Orbitrap Fusion Lumos or QExactive mass spectrometer (Thermo Fisher Scientific inc.).

Raw data were analyzed using Proteome Discoverer (v2.4, Lumos) or Protein Prospector (v5.22.1, QExactive) software against the *E. coli* proteome (https://www.uniprot.org). Search parameters include a precursor and a fragment mass tolerance of 15 ppm and 0.8 Da (Lumos) or 10 ppm and 0.01 Da (QExactive), respectively. Searches included up to two missed trypsin cleavages, and charge states of 2, 3 and 4. Static modification was set to carbamidomethylation of cysteine and dynamic modifications include oxidation of methonines, deamidated glutamine and asparagine, and acetylated protein N-termini. Identified peptides were searched against a random decoy protein database to evaluate the false-positive rate (set to 1%). All raw data is deposited and publicly available in the MassIVE repository (MSV000097236). Gene ontology was performed using https://metascape.org against the *E. coli* proteome using: GO biological processes, KEGG pathway, Reactome Gene Sets, CORUM, TRRUST, PaGenBase and Wiki Pathways.

### Hemagglutination assay

*E. coli* strains were grown statically at 37 °C until OD_600_=1.0; for strains harboring an inducible plasmid, cells were grown till OD_600_=0.6, the gene was induced and grown for another 2 hours. The OD_600_ then was adjusted to 1.0 for comparison between different strains. Guinea pig erythrocytes (Innovative Research, Inc, USA) were washed with PBS and diluted to the final concentration of 3% (v/v). In a 96-well round bottom plate, 50 µL of bacterial suspension was mixed with 50 µL of 3% red blood cell (RBC) suspension. The plate was incubated at room temperature for 60 min without agitation. Absence of type 1 pili is indicated by a “button” formed by the settled RBCs at the round bottom well, whereas presence of type 1 pili binds the RBCs forming a diffuse mixture. For mannose-sensitive hemagglutination, 2% mannose was added to bacterial suspension to selected wells before adding RBC and incubated for 15 min to inhibit type 1 pili-mediated hemagglutination.

### FimA cleavage time point assay

Periplasmic proteins were dialyzed in GlpG activity buffer containing 50 mM MES, pH 6.0, 150 mM NaCl, 20% glycerol, 0.1% DDM, as determined by our group (Arutyunova et al., 2017b). The periplasmic proteins were incubated with 5 µM of *E. coli* GlpG or the catalytically inactive mutant (Ser201Ala) at 37 °C for 6 hr, taking fractions at various time points for western blot analysis.

### Western Blot

To measure the level of FimA over time, fractions of FimA incubated with GlpG were analyzed by standard western blot methods and probed with rabbit anti-FimA antibody (dilution = 1:2,000; Biomatik, CAC11912) and HRP conjugated goat anti-rabbit antibody (dilution = 1:5,000; Thermo Scientific, 31460). Amersham Western Blot Detection Reagent (Cytiva, Cat. #45-002-401) was used and the blot was imaged on the Li-COR using the ImageJ program.

### *In vitro* cleavage assay

FimA FRET peptide, DABCYL-QARYFATGAATPGAANADATFKVQYQ-Glu(EDANS) (BioMatik) was incubated either GlpG in DDM or in PLS. For GlpG in DDM, activity buffer consisting of 50 mM MES, pH 6.0, 150 mM NaCl, 20% glycerol, 0.1% DDM. The reaction was started with 0.5 µM of purified GlpG solubilized in DDM and was monitored for 3 hr at 37 °C. The activity assay with GlpG reconstituted in PLS was performed the same way as for the protease in DDM with the only difference being the activity buffer, where DDM was omitted (50 mM MES pH 6.0, 150 mM NaCl, 20% glycerol). Prism 9.0, GraphPad Software, was used to determine cleavage of the peptide.

### Construction of mutant strains

Ser201Ala point mutations in the *glpG* gene were generated in *E. coli* K12 substr. MG1655 and NU149, denoted as *glpG*-S201A, by method of CRISPR/Cas9 gene-editing described by previous groups (Jiang et al., 2015). Briefly, electrocompetent cells harboring pCas (Addgene: #62225) were co-transformed with pTargetGlpG and 500 ng of donor template DNA containing the *glpG* gene with a S201A mutation. Cells were recovered at 30°C for 2 hr and spread on LB agar containing kanamycin (50 µg/mL) and spectinomycin (50 µg/mL) and incubated overnight at 30°C. Transformants were identified by colony PCR and Sanger sequencing.

### Negatively stained transmission electron microscopy of *E. coli*

Different strains of *E. coli* were grown statically to preserve pili integrity in LB for 48 hr at 37 °C. Strains harboring a GlpG plasmid were induced after the first 8 hours of growth. 7 µL of cell culture were gently added to glow-discharged carbon-coated copper grids (Electron Microscopy Sciences, Cat. # CF200-Cu-50) and left at room temperature for 5 min. Excess sample was wicked away using Wattman paper, washed with nuclease-free water and stained with 2% (w/v) uranyl acetate. Grids were imaged on a Hitachi H-7650 Transmission Electron Microscope.

### RT-qPCR of *glpG*

*E. coli* strains were grown in LB media, TB media, or M9 minimal media to an OD600 of 1.0 and centrifuged at 5000 x *g*. RNA was extracted using TRIzol™ Max Bacterial RNA Isolation kit (Invitrogen, Cat. #16096020). cDNA was amplified using iScript Reverse Transcription Supermix (Bio-RAD, Cat. #1708840) and RT-qPCR was performed with PowerUp™ SYBR™ Green Master Mix (Thermo Scientific, Cat. #A25741) and analyzed for mRNA gene expression using the QuantStudio5 system. Both *glpG* and housekeeping gene *rrsB* were measured with *glpG* mRNA expression was measured as ΔΔCt.

### Mammalian cell culture conditions

Mouse inner medullary collecting duct (mIMCD-3; ATCC, Cat. # CRL-2123) cells were grown in DMEM:F12 medium (Gibco, #Cat. #11320033) containing 10% fetal bovine serum (Sigma-Aldrich, Cat. #12103C) at 37 °C, 5% CO_2_. Human bladder (T24; ATCC, Cat. # HTB-4) cells were grown in McCoy’s medium (Sigma-Aldrich, Cat. #30-2007) containing 10% fetal bovine serum at 37 °C, 5% CO_2_.

### Gentamicin protection assay

Various strains of *E. coli* were grown to an OD_600_ of 1.0 and added to a confluent layer of mouse inner medullary collecting duct (mIMCD-3; ATCC, Cat. # CRL-2123) or human bladder (T24; ATCC, Cat. # HTB-4) cell line to facilitate infection for 4 hr at 37 °C. Cells were washed with sterile PBS and grown in medium containing 100 μg/mL of gentamicin for 2 hr to kill all remaining extracellular bacteria, leaving invasive bacteria unaffected. Invasive bacteria were collected by lysing mammalian cells with 2% (v:v) TritonX-100. Serial dilutions of lysate were plated on LB agar plates and grown for 18 hr at 37 °C to quantify bacterial invasion by manual counting of colony forming units.

### Generation of kidney organoids

Kidney organoids were differentiated using the Takasato protocol with minor modifications as previously described by the Chun group (Chun et al., 2022; Rahmani et al., 2022). Briefly, induced pluripotent stem cells were grown in 6 well plates coated with Matrigel matrix (Corning, Cat. #354277). Cells were treated with 8 μM CHIR99021 (R&D Systems, Cat. #4423) for 4 days followed by recombinant human FGF-9 (200 ng/mL; R&D Systems, Cat. #273-F9-025) and heparin (1 μg/mL, Millipore Sigma, Cat. #H4784) for an additional 3 days in APEL2 medium (Stem Cell Technologies, Cat. #05275) supplemented with 5% PFHM-II (Thermo Fisher, Cat. #12040077). On day 7, cells were dissociated using trypsin-EDTA (0.05%) for 2 min and 100,000 cells were transferred onto a six-well transwell plate (Corning, Cat. # 07-200-170) with nine pellets per well without an aggregation step. Cell clusters were incubated with a pulse of 5 μM CHIR99021 in APEL2 + PFHM-II medium for 1 hr at 37 °C. After 1 hr, the medium was changed to APEL2 + PFHM supplemented with 200 ng/mL FGF-9 and 1 μg/mL heparin for an additional 5 days and then were maintained in APEL2 + PFHM-II medium for 14 days with medium change every other day.

### Infecting kidney organoids with *E. coli*

UPEC *E. coli* NU149 wild-type and NU149 *glpG*-S201A were grown overnight in 5 ml LB at 37 °C, 225 RPM. 300 μl of overnight culture for each strain were transferred to 2.7 ml of LB and grown for 4 hr at 37 °C, 225 RPM. Cultures were centrifuged at 2000 x *g* for 2 min with the supernatant aspirated and discarded. Each pellet of bacteria was resuspended in 1 mL APEL2. On day 26 of differentiation, kidney organoids were infected in triplicates with 1:10 and 1:100 concentrations of *E. coli* strain or vehicle control and infected at 37 °C, 5% CO_2_ for 6 hr with agitation every 1 hr. After 6 hr, supernatant was discarded, and kidney organoids were washed 2 times with PBS for 5 min with rocking for each wash. To inactivate *E. coli*, kidney organoids were treated with an APEL2 solution containing 2X Pen/Strep (200 units/mL penicillin and 200 ug/mL streptomycin) (Gibco; Cat. #15140122) at 37 °C, 5% CO_2_ for 2 hr. The organoids were then washed 2 times with sterile PBS for 5 min and fixed with 2% paraformaldehyde for 20 min at 4 °C. After two washes with PBS, kidney organoids were stored at 4 °C until processed for immunofluorescence and imaging.

### Immunofluorescence and confocal microscopy of kidney organoids

Kidney organoids were processed for immunofluorescence as previously described (Rahmani et al., 2022). In brief, kidney organoids were blocked in a blocking buffer (10% donkey serum and 0.05% Triton-X in PBS) for 2 hr at room temperature. Primary antibodies to *E. coli* (anti-*E. coli*-FITC; Novus Cat. #NB100 64448, 1:25), anti-NPHS1 (R&D Systems, Cat. #AF4269, 1:200), Lotus tetragonolobus lectin-Cy5 (GlycoMatrix Cat. # 21761117-1; 1:200) were incubated at 4 °C overnight. Following the overnight incubation, organoids were washed 3 times with wash buffer (0.3% Triton X-100 in PBS) for 10 min each wash followed by incubation in wash buffer with fluorescent conjugated secondary antibodies that include Alexa Fluor donkey anti-sheep 594 (Thermo Fisher Scientific, 1:250) for 2 hr at room temperature. After 3 washes with PBS and 10 min labeling with 5 μg/ml Hoescht 33342 (ThermoScientific Cat. #H3570), confocal images were acquired with a Zeiss LSM 880 confocal microscope.

### Quantification of bacteria infected proximal tubules

LTL-Cy5 labeled proximal tubules were identified and isolated for quantification of *E. coli* infected proximal tubules of the kidney organoids. Confocal images of *E. coli*-FITC positive structures within the LTL-Cy5 positive regions were analyzed using ImageJ as were described previously for SARS-CoV-2 infected kidney organoids (Chung et al., 2022).

### GlpG inhibition assay

Inhibition was calculated by using 0.2 µM of purified GlpG that was incubated with tosyl phenylalanyl chloromethylketone (TPCK) in a concentration range of 0.1 µM to 5 mM at 37 °C for 60 min in activity buffer consisting of 50 mM MES, pH 6.0, 150 mM NaCl, 20% glycerol, 0.1% DDM. The reaction was started with 15 µM of modified TatA-FRET substrate (K_M_ for modified TatA-FRET is 10 µM; Mca-FATA*AFGSP-Dpn, Biomatik, 98% purity) and was monitored for 3 hr at 37 °C. Prism 9.0, GraphPad Software, was used to determine the IC_50_ value.

### Biofilm formation assay on biotic surface

mIMCD-3 cells were grown in Dulbecco’s Modified Eagle Medium/Nutrient Mixture F-12 (DMEM/F12 1:1, Gibco). Cells were seeded on coverslips pre-coated with poly-L-lysine and grown to 90% confluency. UPEC NU149 strains were grown overnight at 37°C in lysogeny broth (LB). The following day, cells were subcultured in LB at 37°C until the OD_600_ reached 0.2. Protein expression was induced by adding 0.002% arabinose or 1 mM IPTG for 2 h. mIMCD-3 cells were infected with NU149 strains at a multiplicity of infection (MOI) of 1:25 for 1.5 h. Non-adherent bacteria were washed off with 1X PBS twice, then cells were incubated with fresh media for an additional 4 h. At the end of the infection period, cells were washed twice with 1X PBS, then fixed with 3% paraformaldehyde (Sigma-Aldrich) for 30 min at room temperature. Coverslips were blocked overnight with 3% goat serum (Invitrogen) and 1% bovine serum albumin in PBS (BioShop) at 4°C. To stain the cells and bacteria, samples were incubated with rabbit anti-O-Ag conjugated to FITC (1:50 dilution, Novus Biologicals) and phalloidin Alexa Fluor 568 (1:40 dilution, Invitrogen, CAT: A12380) for 1 h at room temperature. Coverslips were washed three times in 1x PBST, then stained with 300 nM DAPI (Invitrogen, CAT: D1306) for 5 min at room temperature and washed three times with 1x PBST prior to mounting on glass slides. All images were captured with BioTek Cytation 5 Imager using a 20X objective. Samples were imaged blindly and processed with BioTek Gen5 software for biofilm quantification. Acquired images were preprocessed then analyzed to quantify aggregates with surface area of 100 µm^2^ or more, shown previously to represent biofilms formed by pathogenic *E. coli* (CITE Elhenawy et al 2021). Images were obtained from three independent experiments.

### Biofilm formation assay on abiotic surface

*E. coli* NU149 wild-type, *E. coli* NU149 *glpG*-S201A strain and *E. coli* NU149 *glpG-*S201A + GlpG, the inactive *glpG* strain transformed with pBad plasmid bearing active GlpG protease, were streaked on LB agar with the appropriate antibiotic. Bacteria cultures were grown in LB medium from colonies and grown for 16-18 hr at 37 °C with shaking. Overnight cultures were diluted to OD_600_ of 0.01 where 200 µl of bacterial sample was added per well on a 96-well plate. LB media alone was always included as a negative control of contamination. Bacterial cultures were grown at 37 °C under static conditions. After 3 hr, arabinose was added to the wells with NU149 *glpG-* S201A + GlpG strain to the final concentration of 0.02%. After 24 hr, the optical density at 600 nm was recorded on a BioTeK Cytation5 microplate reader to quantify the overall bacterial growth in each well. Media with planktonic cells was discarded and wells were rinsed three times with PBS. Once excess PBS was removed, adhered biomass was stained by adding 230 µl of 0.1% Safranin solution to each well and placing the plate on an orbital shaker for 30 min at room temperature (Ommen et al., 2017). The stain was discarded, the wells were rinsed with PBS three times again and the excess of the buffer was removed. Safranin dye bound to the adhered biofilm was solubilized in 230 µl of 30% acetic acid per well. The absorbance at 595 nm was recorded to evaluate the amount of adhered biomass.

### Biofilm inhibition assay

TPCK inhibitor was added at the same time as a freshly prepared bacterial suspension with OD_600_ of 0.01 at the final concentration of 400 µM and then incubating for a certain time followed by quantification of biofilm biomass by Safranin the same way as described above.

### Biofilm eradication assay

A requirement is pre-establishing consistent biofilms in the well prior to inhibition, thus biofilm was allowed to be formed as described above. After incubation for 24 hr the adhered biomass was rinsed with fresh LB media, treated with TPCK (400 µM final concentration), incubated for another 24 hr and quantified with Safranin dye.

### Metabolic activity assay

The 96 well plate for metabolic activity assay was set up the same way as the biofilm eradication assay plate above. After pre-establishing biofilms, washed with fresh LB media and treated with TPCK inhibitor, TTC dye (2,3,5-triphenyl tetrazolium chloride, 0.05% final concentration) was added to the wells and the plate was incubated for 24 hr under static conditions at 37 °C. The wells were rinsed three times with PBS, and 230 µl of methanol was added to each well followed by quantification by reading the absorbance at 500 nm.

### LC-MS/MS proteomics on whole cells

*E. coli* strain DY330 harboring a triple FLAG-tag inserted at the N-terminus of the genomic *glpG* gene was grown in LB media with kanamycin and lysed in buffer containing 50 mM Tris, pH 8.0, 200 mM NaCl, 20% glycerol, 1% Triton X-100. Cleared fractions were examined using LC-MS/MS. The Orbitrap Elite mass spectrometer was operated with Thermo XCalibur software. All RAW files were converted to mzXML using ReAdW-4.3.1 and submitted for database searching using SEQUEST-PVM v.27 (rev. 9) (Eng et al., 1994) under standard workflow conditions with a non-redundant E. coli proteome sequence FASTA file from Ecocyc. Search parameters were set to allow for dynamic modification of methionine oxidation and fixed modification of cysteine carbamidomethylation using precursor ion tolerance of 20 ppm. A stringent false discovery rate of 1% (P 0.01) was used to filter candidate peptide and protein identifications. Precursor mass tolerance was set to 3 Da (daughter mass ion tolerance set to the default of 0), while enabling partial tryptic enzyme and single site missed cleavages. The STATQUEST filtering algorithm (Kislinger et al., 2003) was then applied to all putative SEQUEST search results to assign statistical confidence. High confidence (>90% likelihood) spectral counts from SEQUEST/STATQUEST were used to calculate NSAF score (Florens et al., 2006). To be more conservative, the proteins were then filtered on median NSAF score or STATQUEST probability of 99%. Proteins localized in the periplasm and inner membrane (Babu et al., 2018), belonging to the COG categories O and U (Tatusov et al., 1997) were visualized as Cytoscape network (Shannon et al., 2003). All raw data is deposited and publicly available in the MassIVE repository (MSV000097249).

## Supporting information

Data Set 1

Data Set 2

Data Set 3

## Acknowledgements

We thank Jack Moore and the Alberta Proteomics and Mass Spectrometry Facility for technical support. We also thank Sara Amidian, Joesph Primeau for microscopy support at the University of Alberta Imaging Core, and the SPP-ARC Cryo-EM Facility. We thank Dr. Tracy Raivio for supplying pathogenic *E. coli* and K12 single gene knockouts. We received funding from the Canadian Foundation for Innovation (37833 and 39051) and Natural Sciences, Engineering Research Council of Canada (DGECR-2018-00142 discovery grant to O.J.) and Alberta Graduate Excellence Scholarship (B.H).

This work was supported by the Women’s and Children’s Health Research Institute at the University of Alberta (MJL), the National Science and Engineering Research Council of Canada (RGPIN-2016-06478 to MJL; RGPIN-2019-20234 to MB) and RGPIN-2023-04396 (MJL), Striving for Pandemic Preparedness – The Alberta Research Consortium, and Canada Foundation for Innovation (CFI), Canada Research Chair CRC-2021-00172 (HKA), Canadian Institutes of Health Research (PJT-180387 to HSY; PJT-426160 to EW).

For experiments performed at the University of Alberta Faculty of Medicine & Dentistry Cell Imaging Core, RRID:SCR_019200, which receives financial support from the Faculty of Medicine & Dentistry, the University Hospital Foundation, Striving for Pandemic Preparedness – The Alberta Research Consortium, and Canada Foundation for Innovation (CFI) awards to contributing investigators.

## Author Contributions

Conceptualization, J.L., E.A. and M.J.L.; Affinity-purification-mass spectrometry, S.P and M.B., In-gel digestion proteomics, B.H. and O.J.; Generation of mutant *E. coli*, protein purification, immunoblotting, sample preparation for electron microscopy, RT-qPCR, J.L., Gentamicin protection assay, J.L., L.B., H.A. and E.W., Abiotic biofilm assays, inhibition assays, metabolic assays, enzyme kinetics, E.A., Kidney organoid generation and infection, microscopy of organoids, H.J.C. and J. C., Biotic biofilm assays, J.W. and W.E., Writing – Original Draft, J.L. and E.A.; Writing – Review & Editing, J.L., E.A. and M.J.L.; Supervision, M.J.L., H.S.Y., O.J., J. C., W.E and M.B.; Funding Acquisition, M.J.L., H.S.Y., O.J., J. C., W.E and M.B.

## Inclusion and Diversity

We support inclusive, diverse, and equitable conduct of research

