## Supplemental Figures 1-10 for "Mechanistic insight into the regulation of virulence factor type 1 pili in pathogenic *E. coli* by rhomboid protease GlpG"

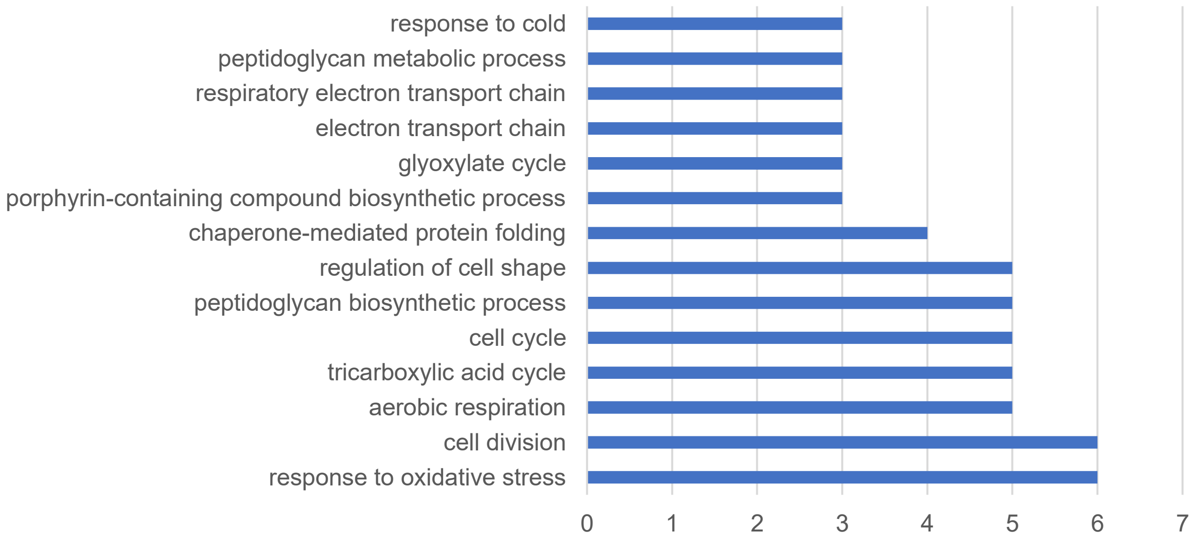


**Supplemental Figure 1.** Gene ontology groups of decreased periplasmic proteins when active GlpG is added, identified in LC-MS/MS

| Accession | UniProt ID | Gene name | Description | Abundance Ratio (log2): | (-)LOG(P-value) |
|---|---|---|---|---|---|
| P0ADB7 | ECNB | ecnB | Entericidin B | -6.64 | 6 |
| P16676 | CYSA | cysA | Sulfate/thiosulfate import ATP-binding protein CysA | -6.64 | 6 |
| P76027 | OPPD | oppD | Oligopeptide transport ATP-binding protein OppD | -6.64 | 6 |
| P64534 | RCNB | rcnB | Nickel/cobalt homeostasis protein RcnB | -6.64 | 6 |
| P0AFM6 | PSPA | pspA | Phage shock protein A | -6.64 | 6 |
| P0AFM9 | PSPB | pspB | Phage shock protein B | -6.64 | 6 |
| P10346 | GLNQ | glnQ | Glutamine transport ATP-binding protein GlnQ | -6.64 | 6 |
| P0AAE0 | CYCA | cycA | D-serine/D-alanine/glycine transporter | -6.64 | 6 |
| P0AFF2 | NUPC | nupC | Nucleoside permease NupC | -6.64 | 6 |
| P32705 | ACTP | actP | Cation/acetate symporter ActP | -6.64 | 6 |
| P69741 | MBHT | hybO | Hydrogenase-2 small chain | -6.64 | 6 |
| P0AEH5 | ELAB | elaB | Protein ElaB | -6.64 | 6 |
| P27294 | INAA | inaA | Protein InaA | -5.43 | 3.336299075 |
| P0AGI8 | TRKA | trkA | Trk system potassium uptake protein TrkA | -5.13 | 5.22184875 |
| P77757 | ARNC | arnC | Undecaprenyl-phosphate 4-deoxy-4-formamido-L-arabinose transferase | -4.85 | 3.110138279 |
| P0AFD6 | NUOI | nuoI | NADH-quinone oxidoreductase subunit I | -4.84 | 6 |
| P00393 | NDH | ndh | Type II NADH:quinone oxidoreductase | -4.64 | 4.229147988 |
| P0ADW3 | YHCB | yhcB | Inner membrane protein YhcB | -4.43 | 3.022733788 |
| P0A754 | KEFF | kefF | Glutathione-regulated potassium-efflux system ancillary protein KefF | -4.28 | 3.41453927 |
| P0A9R7 | FTSE | ftsE | Cell division ATP-binding protein FtsE | -4.25 | 2.780677492 |
| P0ABH0 | FTSA | ftsA | Cell division protein FtsA | -4 | 6 |
| P25746 | HFLD | hflD | High frequency lysogenization protein HflD | -3.99 | 3.795880017 |
| P24228 | DACB | dacB | D-alanyl-D-alanine carboxypeptidase DacB | -3.98 | 6 |
| P0ACB4 | HEMG | hemG | Protoporphyrinogen IX dehydrogenase [quinone] | -3.85 | 3.920818754 |
| P07000 | PLDB | pldB | Lysophospholipase L2 | -3.58 | 2.81276138 |
| P23894 | HTPX | htpX | Protease HtpX | -3.42 | 2.103473783 |
| P0A996 | GLPC | glpC | Anaerobic glycerol-3-phosphate dehydrogenase subunit C | -3.11 | 2.361710465 |
| P0AC41 | SDHA | sdhA | Succinate dehydrogenase flavoprotein subunit | -2.9 | 5 |
| P77302 | DGCM | dgcM | Diguanylate cyclase DgcM | -2.81 | 4.958607315 |
| P21513 | RNE | rne | Ribonuclease E | -2.75 | 5.522878745 |
| P0A6J5 | DADA | dadA | D-amino acid dehydrogenase | -2.75 | 4.886056648 |
| P06149 | DLD | dld | Quinone-dependent D-lactate dehydrogenase | -2.52 | 2.399462706 |
| P0AFC7 | NUOB | nuoB | NADH-quinone oxidoreductase subunit B | -2.52 | 2.301290651 |
| P0A9X1 | ZNUC | znuC | Zinc import ATP-binding protein ZnuC | -2.48 | 4.366531544 |
| P37690 | ENVC | envC | Murein hydrolase activator EnvC | -2.46 | 2.30565809 |
| P69874 | POTA | potA | Spermidine/putrescine import ATP-binding protein PotA | -2.33 | 1.709408957 |
| P0AFD1 | NUOE | nuoE | NADH-quinone oxidoreductase subunit E | -2.17 | 1.552407891 |
| P04128 | FIMA1 | fimA | Type-1 fimbrial protein, A chain | -2.16 | 3.208309351 |
| P0AAF3 | ARAG | araG | Arabinose import ATP-binding protein AraG | -2.07 | 3.056505484 |
| P0ABB0 | ATPA | atpA | ATP synthase subunit alpha | -2.05 | 3.18243463 |
| P23830 | PSS | pssA | CDP-diacylglycerol--serine O-phosphatidyltransferase | -2.02 | 2.545917729 |
| P07014 | SDHB | sdhB | Succinate dehydrogenase iron-sulfur subunit | -1.83 | 1.400717499 |
| P0A8C1 | YBJQ | ybjQ | UPF0145 protein YbjQ | -1.59 | 3.554395797 |
| P0ACB7 | HEMY | hemY | Protein HemY | -1.53 | 2.27392513 |
| P0AAG8 | MGLA | mglA | Galactose/methyl galactoside import ATP-binding protein MglA | -1.47 | 3.059483515 |
| P0AEB2 | DACA | dacA | D-alanyl-D-alanine carboxypeptidase DacA | -1.45 | 3.661543506 |
| P0ADT8 | YGIM | ygiM | Uncharacterized protein YgiM | -1.39 | 2.228339041 |
| P31979 | NUOF | nuoF | NADH-quinone oxidoreductase subunit F | -1.2 | 2.370488466 |
| P0A7J0 | RIBB | ribB | 3,4-dihydroxy-2-butanone 4-phosphate synthase | -1.19 | 3.009217308 |
| P0A901 | BLC | blc | Outer membrane lipoprotein Blc | -1.13 | 1.835141753 |

**Supplemental Figure 2.** Periplasmic proteins identified as significantly decreased in the presence of active GlpG by factor of log-2 when analysed with LC-MS/MS. Periplasmic proteins were isolated from K12 substr. MG1655 *glpG*-KO mutant. Full data set is in **Data S1**

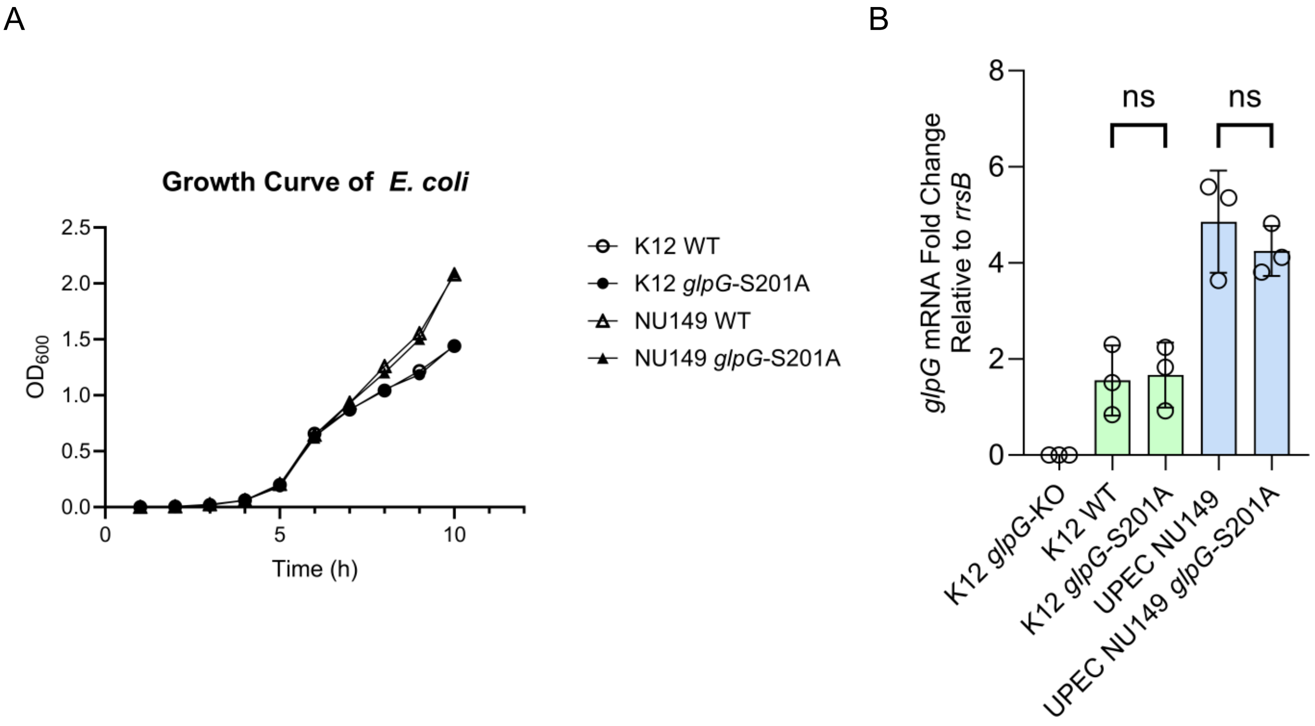


**Supplemental Figure 3.** A) Growth assay comparing K12 and NU149 WT to their respective *glpG*-S201A genomic mutant. B) Changes in *glpG* mRNA levels are not significant when the *glpG* gene is endogenously mutated to its inactive form (Ser201Ala) using CRISPR/Cas. Significant differences were determined by one-way ANOVA.

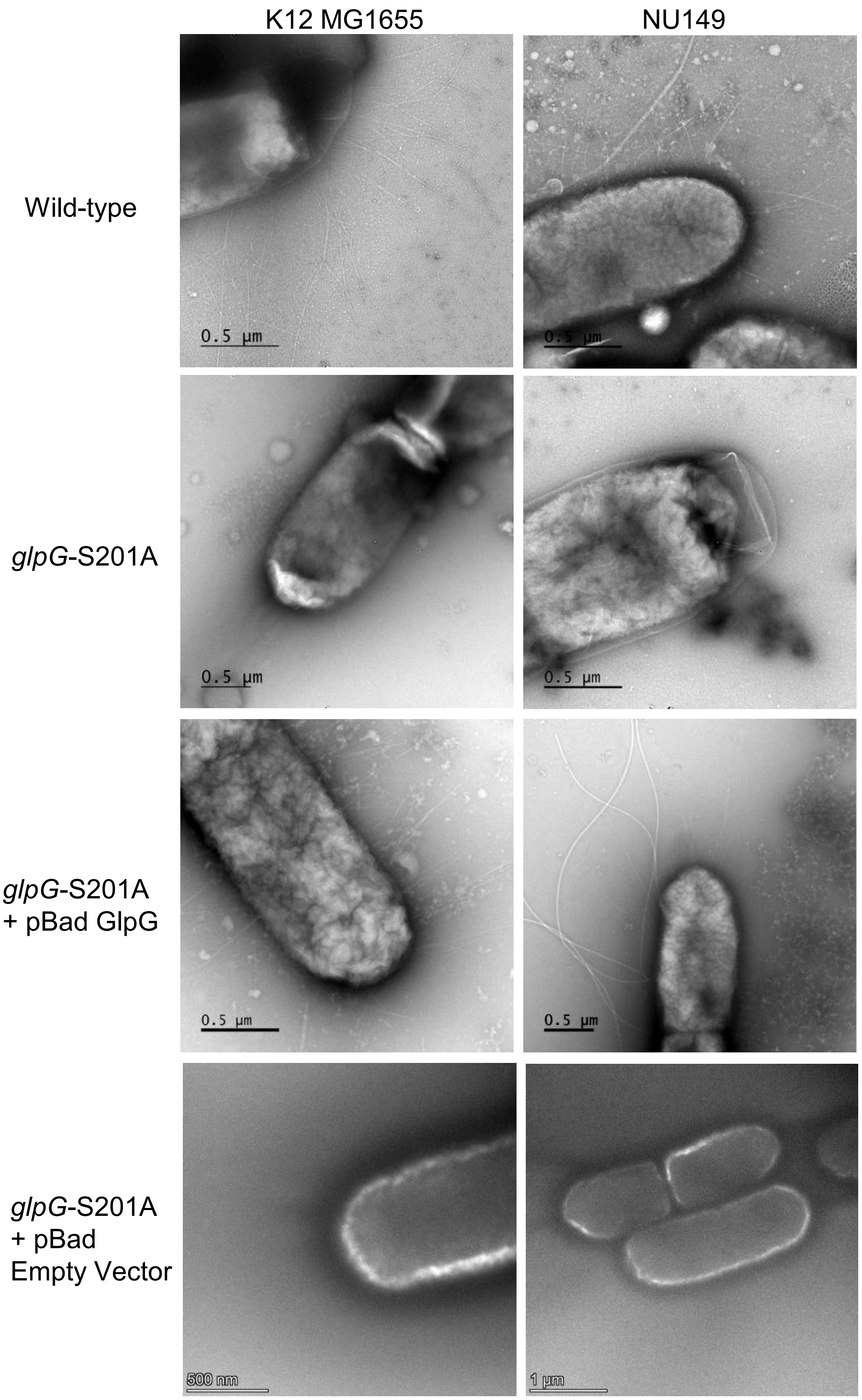


**Supplemental Figure 4.** Surface pili formation is impaired when GlpG activity is not present in both laboratory and uropathogenic *E. coli* strains. Left: K12 MG1655 wild-type, CRISPR/Cas9 mutant, CRISPR/Cas9 mutant overexpressing active GlpG in a pBad inducible plasmid, and CRISPR/Cas9 harboring an empty pBad plasmid. Right: UPEC strain NU149 wild-type, CRISPR/Cas9 mutant, CRISPR/Cas9 mutant overexpressing active GlpG in a pBad inducible plasmid, and CRISPR/Cas9 harboring an empty pBad plasmid.

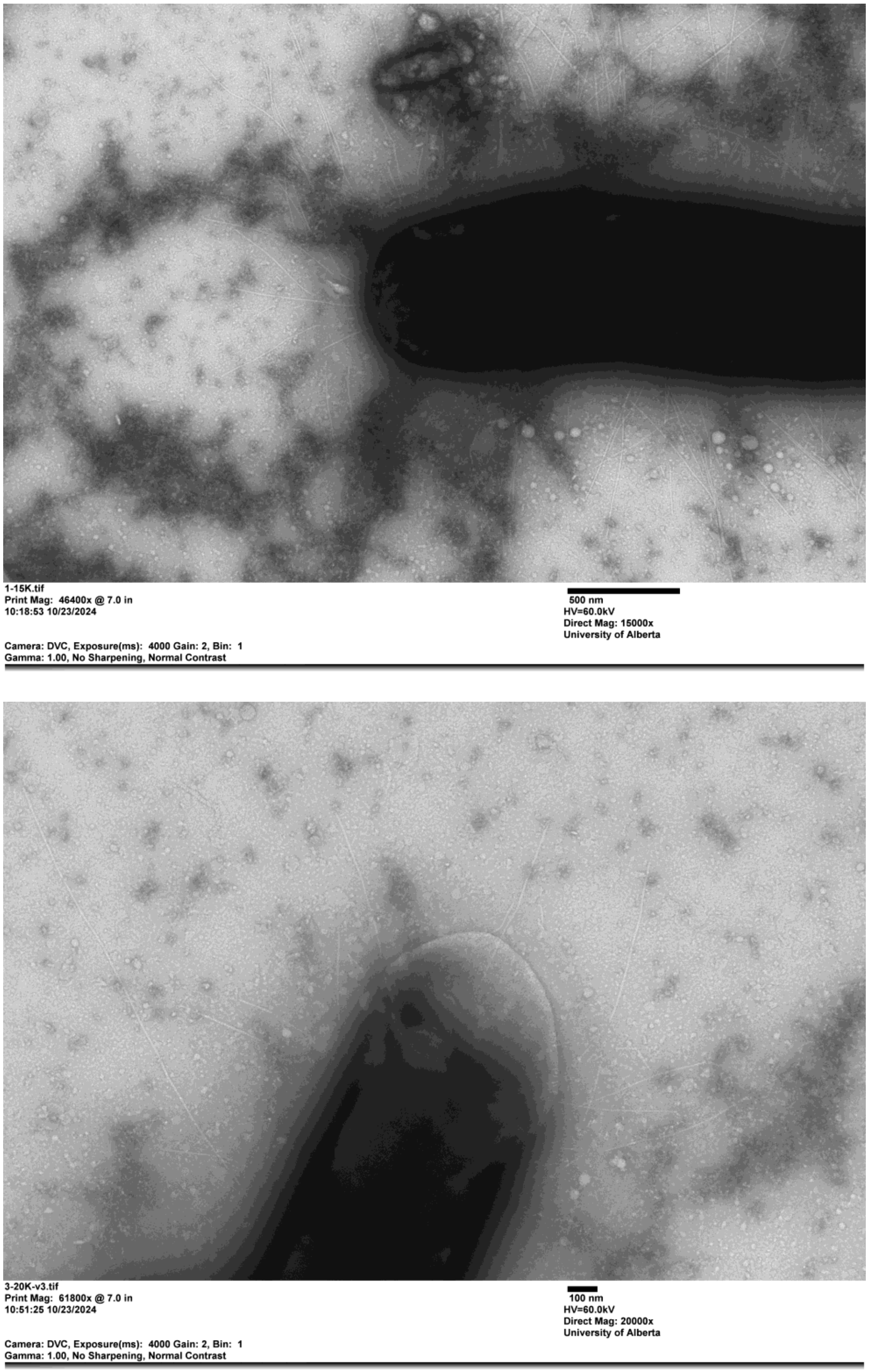


**Supplemental Figure 5.** NU149 *glpG*-S201A overexpressing FimC using an inducible plasmid shows rescue in pili formation

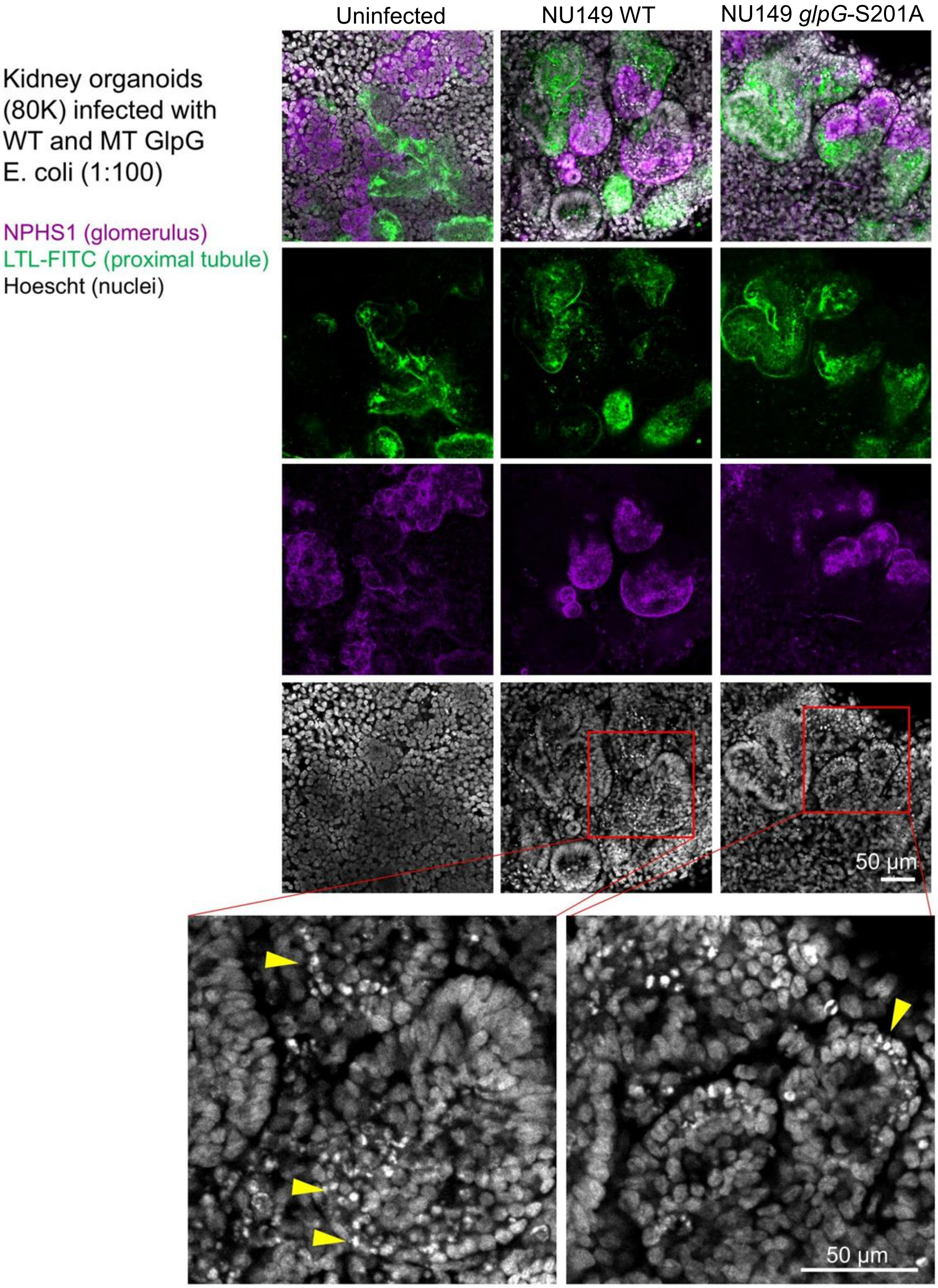


**Supplemental Figure 6** Kidney organoids indicating the development of proximal tubules and visualization of *E. coli* invasion, represented by small puncta, depicted by yellow arrows.

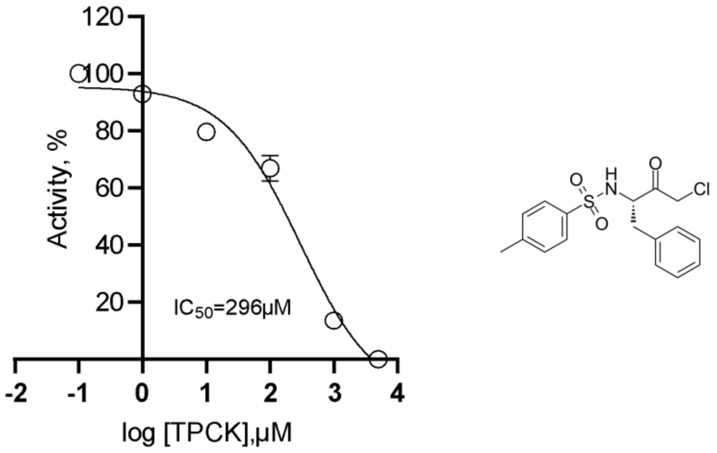


**Supplemental Figure 7.** Dose response curve and corresponding $IC_{50}$ value of TPCK for *E. coli* GlpG protease.

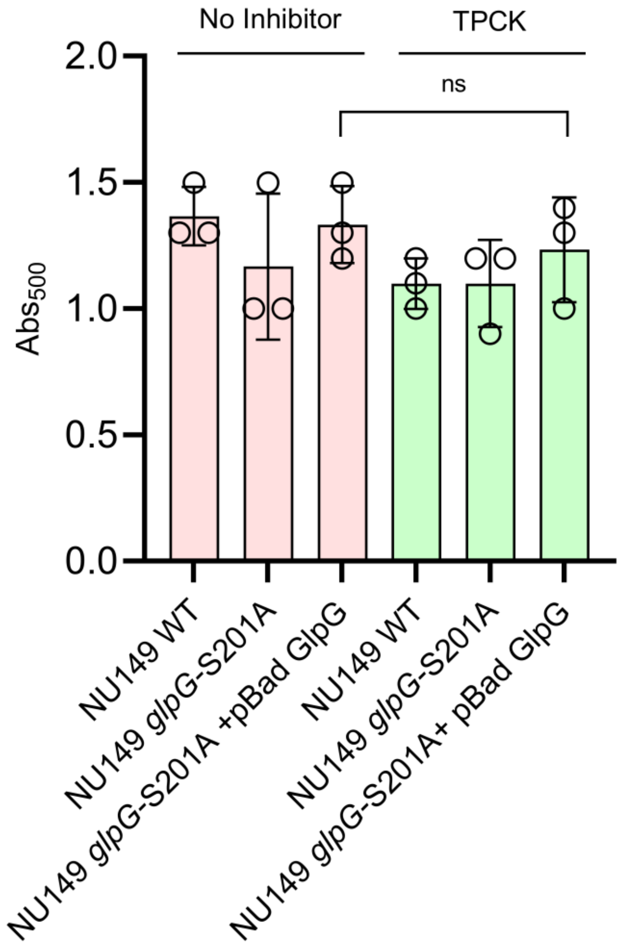


**Supplemental Figure 8.** Absorbance at 500 nm of solubilized TTC dye quantifying the metabolic activity of bacteria in the presence and absence of TPCK. Significant changes were determined by one-way ANOVA. ns: $P>0.05$.

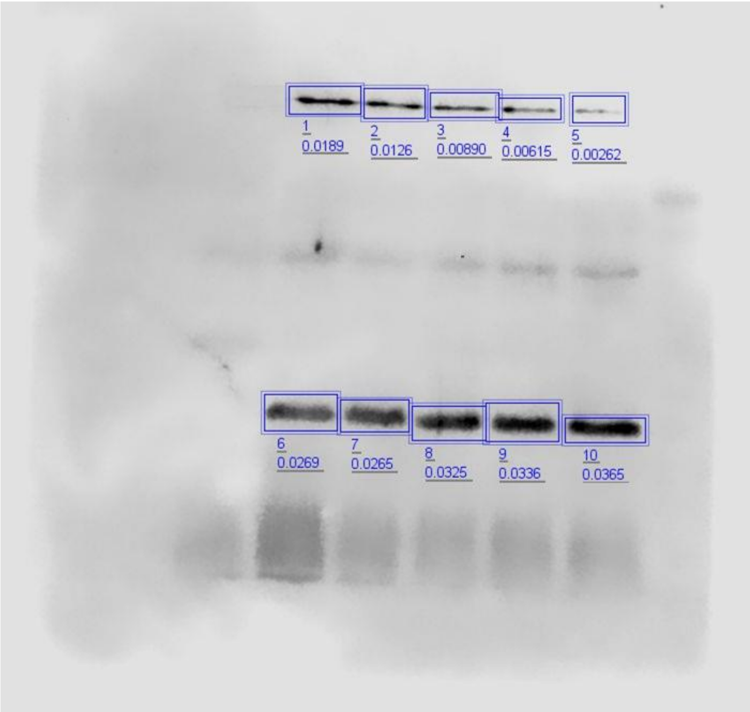


**Supplemental Figure 9.** Densitometry analysis quantifying FimA cleavage over time in the presence of active DDM-solubilized GlpG
